# The neural representation of force across grasp types in motor cortex of humans with tetraplegia

**DOI:** 10.1101/2020.06.01.126755

**Authors:** Anisha Rastogi, Francis R. Willett, Jessica Abreu, Douglas C. Crowder, Brian A. Murphy, William D. Memberg, Carlos E. Vargas-Irwin, Jonathan P. Miller, Jennifer Sweet, Benjamin L. Walter, Paymon G. Rezaii, Sergey D. Stavisky, Leigh R. Hochberg, Krishna V. Shenoy, Jaimie M. Henderson, Robert F. Kirsch, A. Bolu Ajiboye

## Abstract

Intracortical brain-computer interfaces (iBCIs) have the potential to restore hand grasping and object interaction to individuals with tetraplegia. Optimal grasping and object interaction require simultaneous production of both force and grasp outputs. However, since overlapping neural populations are modulated by both parameters, grasp type could affect how well forces are decoded from motor cortex in a closed-loop force iBCI. Therefore, this work quantified the neural representation and offline decoding performance of discrete hand grasps and force levels in two participants with tetraplegia. Participants attempted to produce three discrete forces (light, medium, hard) using up to five hand grasp configurations. A two-way Welch ANOVA was implemented on multiunit neural features to assess their modulation to *force* and *grasp*. Demixed principal component analysis was used to assess for population-level tuning to force and grasp and to predict these parameters from neural activity. Three major findings emerged from this work: 1) Force information was neurally represented and could be decoded across multiple hand grasps (and, in one participant, across attempted elbow extension as well); 2) Grasp type affected force representation within multi-unit neural features and offline force classification accuracy; and 3) Grasp was classified more accurately and had greater population-level representation than force. These findings suggest that force and grasp have both independent and interacting representations within cortex, and that incorporating force control into real-time iBCI systems is feasible across multiple hand grasps if the decoder also accounts for grasp type.

**Significance Statement:** Intracortical brain-computer interfaces (iBCIs) have emerged as a promising technology to potentially restore hand grasping and object interaction in people with tetraplegia. This study is among the first to quantify the degree to which hand grasp affects force-related – or *kinetic* – neural activity and decoding performance in individuals with tetraplegia. The study results enhance our overall understanding of how the brain encodes kinetic parameters across varying kinematic behaviors -- and in particular, the degree to which these parameters have independent versus interacting neural representations. Such investigations are a critical first step to incorporating force control into human-operated iBCI systems, which would move the technology towards restoring more functional and naturalistic tasks.

## Introduction

Intracortical brain-computer interfaces (iBCIs) have emerged as a promising technology to restore upper limb function to individuals with paralysis. Traditionally, iBCIs decode kinematic parameters from motor cortex to control the position and velocity of end effectors. These iBCIs evolved from the seminal work of Georgopoulos and colleagues, who proposed that motor cortex encodes high-level kinematics, such as continuous movement directions and three-dimensional hand positions, in a global coordinate frame (Georgopoulos et al., 1982; Georgopoulos et al., 1986). Kinematic iBCIs have successfully achieved control of one- and two-dimensional computer cursors (Wolpaw et al., 2002; Leuthardt et al., 2004; Kubler et al., 2005; Hochberg et al., 2006; Kim et al., 2008; Schalk et al., 2008; Hermes et al., 2011; Kim et al., 2011; Simeral et al., 2011); prosthetic limbs (Hochberg et al., 2012; Collinger et al., 2013; Wodlinger et al., 2015); and paralyzed arm and hand muscles (Bouton et al., 2016; Ajiboye et al., 2017).

While kinematic iBCIs can restore basic reaching and grasping movements, restoring the ability to grasp and interact with objects requires both kinematic and kinetic (force-related) information (Chib et al., 2009; Flint et al., 2014; Casadio et al., 2015). Specifically, sufficient contact force is required to prevent object slippage; however, excessive force may cause mechanical damage to graspable objects (Westling and Johansson, 1984). Therefore, introducing force calibration capabilities during grasp control would enable iBCI users to perform more functional tasks.

Early work by Evarts and others, which showed correlations between cortical activity and force output (Evarts, 1968; Humphrey, 1970; Fetz and Cheney, 1980; Evarts et al., 1983; Kalaska et al., 1989), and later work, which directly decoded muscle activations from neurons in primary motor cortex (M1) (Morrow and Miller, 2003; Sergio and Kalaska, 2003; Pohlmeyer et al., 2007; Oby et al., 2010), suggest that cortex encodes low-level dynamics of movement along with kinematics (Kakei et al., 1999; Carmena et al., 2003; Branco et al., 2019). However, explorations of kinetic information as a control signal for iBCIs have only just begun. The majority have characterized neural modulation to executed kinetic tasks in primates and able-bodied humans (Filimon et al., 2007; Moritz et al., 2008; Pohlmeyer et al., 2009; Ethier et al., 2012; Flint et al., 2012; Flint et al., 2014; Flint et al., 2017; Schwarz et al., 2018). Small subsets of M1 neurons have been used to command muscle activations through FES to restore one-dimensional wrist control and whole-hand grasping in non-human primates with temporary motor paralysis (Moritz et al., 2008; Pohlmeyer et al., 2009; Ethier et al., 2012). More recent intracortical studies demonstrated that force representation is preserved in individuals with chronic tetraplegia (Downey et al., 2018; Rastogi et al., 2020).

Intended forces are usually produced in the context of task-related factors, including grasp postures used to generate forces (Murphy et al., 2016). The representation and decoding of grasps – independent of forces – has been studied extensively in non-human primates (Stark and Abeles, 2007; Stark et al., 2007; Vargas-Irwin et al., 2010; Carpaneto et al., 2011; Townsend et al., 2011; Hao et al., 2014; Schaffelhofer et al., 2015) and humans (Pistohl et al., 2012; Chestek et al., 2013; Bleichner et al., 2014; Klaes et al., 2015; Bleichner et al., 2016; Leo et al., 2016; Branco et al., 2017). Importantly, previous studies suggest that force and grasp are encoded by overlapping populations of neural activity (Sergio and Kalaska, 1998; Carmena et al., 2003; Sergio et al., 2005; Sburlea and Muller-Putz, 2018). While some studies suggest that force is encoded at a high level independent of motion and grasp (Chib et al., 2009; Hendrix et al., 2009; Pistohl et al., 2012; Intveld et al., 2018), others suggest that it is encoded at a low level intertwined with grasp (Hepp-Reymond et al., 1999; Degenhart et al., 2011). Thus, the degree to which intended hand grasps and forces interact within the neural space, and how such interactions affect force decoding performance, remain unclear. To our knowledge, these scientific questions have not been explored in individuals with tetraplegia, who constitute a target population for iBCI technologies.

To answer these questions, we characterized the extent to which three discrete, attempted forces were neurally represented and offline-decoded across up to five hand grasp configurations in two individuals with tetraplegia. Our results suggest that force has both grasp-independent and grasp-dependent (interacting) representation in motor cortex. Additionally, while this study demonstrates the feasibility of incorporating discrete force control into human-operated iBCIs, these systems will likely need to incorporate grasp and other task parameters to achieve optimal performance.

## Materials and Methods

### Study permissions and participants

Study procedures were approved by the US Food and Drug Administration (Investigational Device Exemption #G090003) and the Institutional Review Boards of University Hospitals Case Medical Center (protocol #04-12-17), Stanford University (protocol #20804), and Massachusetts General Hospital (2011P001036). The present study collected neural recordings from participants enrolled in the BrainGate2 Pilot Clinical Trial (ClinicalTrials.gov number NCT00912041). The current work utilized the recording opportunity afforded by the BrainGate2 Pilot Clinical Trial but reports no clinical trial outcomes or endpoints. Informed consent, including consent to publish, was obtained from the participants prior to their enrollment in the study.

This present study includes data from two participants with chronic tetraplegia. Both participants had two, 96-channel microelectrode intracortical arrays (1.5 mm electrode length, Blackrock Microsystems, Salt Lake City, UT) implanted in the hand and arm area (“hand knob”) (Yousry et al., 1997) of dominant motor cortex. Participant T8 was a 53-year-old right-handed male with C4-level AIS-A spinal cord injury 8 years prior to implant; and T5 was a 63-year-old right-handed male with C4-level AIS-C spinal cord injury. More surgical details can be found at (Ajiboye et al., 2017) for T8 and (Pandarinath et al., 2017) for T5.

### Participant Task

The goal of this study was to measure the degree to which various hand grasps affect decoding of grasp force from motor cortical spiking activity. To this end, participants T8 and T5 took part in several research sessions in which they attempted to produce three discrete forces (light, medium, hard) using one of four designated hand grasps (closed pinch, open pinch, ring pinch, power). T8 completed six sessions between trial days 735-956 relative to the date of his microelectrode array placement surgery; and T5 completed one session on trial day 390. During Session 5, participant T8 completed five additional blocks in which he attempted to produce discrete forces during attempted elbow extension. Table 1 lists all relevant sessions and their associated task parameters.

**Table 1.** Session information. Session information for participants T8 and T5, including the number of blocks per grasp type.

| Session No. | Participant | Post-Implant Day | No. Blocks Per Grasp |  |  |  |  |
| --- | --- | --- | --- | --- | --- | --- | --- |
|  |  |  | Closed Pinch | Open Pinch | Ring Pinch | Power | Elbow |
| 1 | T8 | Day 735 | 11 |  |  | 10 |  |
| 2 | T8 | Day 771 | 5 | 5 | 5 | 5 |  |
| 3 | T8 | Day 774 | 6 | 5 | 5 | 5 |  |
| 4 | T8 | Day 788 | 5 | 5 | 5 | 5 |  |
| 5 | T8 | Day 802 | 4 | 4 | 4 | 4 | 5 |
| 6 | T8 | Day 956 | 4 | 4 | 4 | 4 |  |
| 7 | T5 | Day 390 | 4 | 4 | 4 | 4 |  |

Each research session consisted of multiple 4-minute data collection blocks, which were each assigned to a particular hand grasp, as illustrated in Figure 1B. Blocks were presented in a pseudorandom order, in which hand grasps were assigned randomly to each set of two (Session 1), four (Sessions 2-4, 6-7), or five (Session 5) blocks. This allowed for an equal number of blocks per hand grasp, distributed evenly across the entire research session.

**Figure 1.**
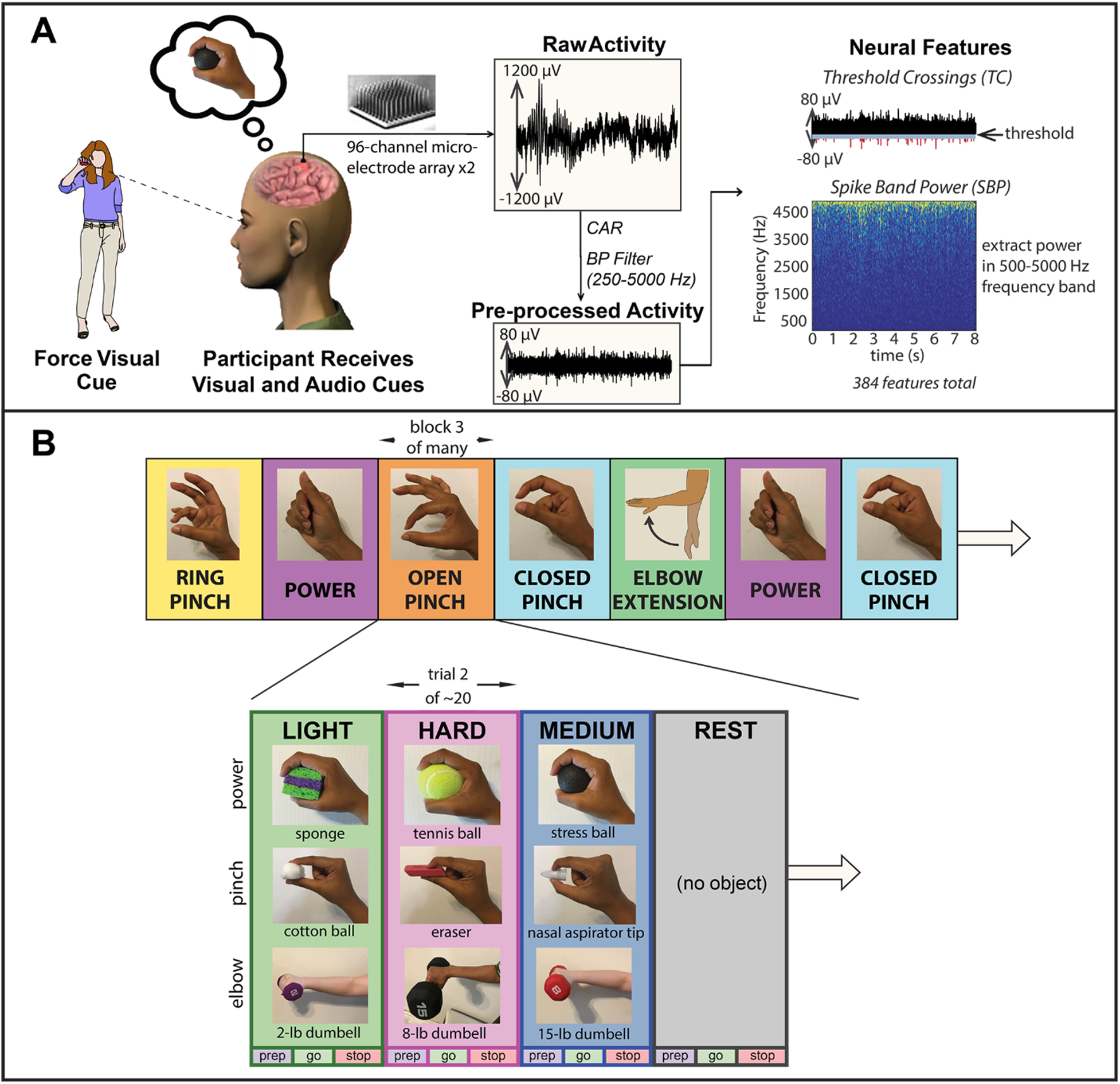
**A.** Experimental setup (Reproduced from (Rastogi et al., 2020)). Participants had two 96-channel microelectrode arrays placed chronically in motor cortex, which recorded neural activity while participants completed a force task. Threshold crossing (TC) and spike band power (SBP) features were extracted from these recordings. **B.** Research session architecture. Each session consisted of 12-21 blocks, each of which contained ~20 trials (see Table 1). In each trial, participants attempted to generate one of three visually-cued forces with one of four grasps: power, closed pincer (c-pinch), open pinch (o-pinch), ring pinch (r-pinch). During session 5, participant T8 also attempted force production using elbow extension. Each trial contained a preparatory (prep) phase, a go phase where forces were actively embodied, and a stop phase where neural activity was allowed to return to baseline. Participants were prompted with both audio and visual cues, in which a researcher squeezed or lifted an object associated with each force level.

All blocks consisted of approximately 20 trials, which were presented in a pseudorandom order by repeatedly cycling through a complete, randomized set of force levels until the end of the block. During each trial, participants used kinesthetic imagery (Stevens, 2005; Mizuguchi et al., 2017) to internally emulate one of three discrete force levels, or rest, with the dominant hand. Participants received simultaneous audio and visual cues indicating which force to produce, when to produce it, and when to relax. Participants were visually cued by observing a researcher squeeze one of nine graspable objects corresponding to light, medium, and hard forces (no object was squeezed during “rest” trials), as shown in Figure 1B. The participants were asked to “follow along” and attempt the same movements that the researcher was demonstrating. The graspable objects were grouped into three sets of three, corresponding to forces embodied using a power grasp (sponge = light, stress ball = medium, tennis ball = hard); a pincer grasp (cotton ball = light, nasal aspirator tip = medium, eraser = hard); or elbow extension (5-lb dumbbell = light, 10-lb dumbbell = medium, 15-lb dumbbell = hard). During the prep phase, which lasted a pseudo-randomly determined period between 2.7 and 3.3 seconds to reduce confounding effects from anticipatory activity, the researcher presented an object indicating the force level to be attempted. The researcher then squeezed the object (or lifted the object, in the case of elbow extension) during the go phase (3-5 seconds), and finally released the object at the beginning of the stop phase (5 seconds).

### Neural Recordings

#### Pre-processing

In both participants, each intracortical microelectrode array was attached to a percutaneous pedestal connector on the head. Cables connected the pedestals to amplifiers (Blackrock Microsystems, Salt Lake City, UT) that bandpass filtered (0.3 Hz – 7.5 kHz) and digitized (30 kHz) the neural signals from each channel on the microelectrode array. These digitized signals were pre-processed in Simulink using the xPC real-time operating system (The Mathworks Inc., Natick, MA, US). Each channel was bandpass filtered (250-5000 Hz), common average referenced (CAR), and down-sampled to 15 kHz in real time. CAR was implemented by selecting 60 channels from each microelectrode array that exhibited the lowest variance, and then averaging these channels together to yield an array-specific common average reference. This reference signal was subtracted from the signals from all channels within each of the arrays.

#### Extraction of Neural Features

From each filtered, CAR channel, two neural features were extracted in real time using the xPC operating system unsorted threshold crossing (TC) and spike band power (SBP) features, from non-overlapping 20 millisecond time bins, as illustrated in Figure 1A. TC features were defined as the number of times the voltage on each channel crossed a predefined noise threshold (Christie et al, 2015) within each time bin (−4.5 × root mean square voltage). Root mean square (RMS) voltage was calculated from one minute of neural data recorded at the beginning of each research session. SBP features were defined as the RMS of the signal in the spike band (250-5000 Hz) of each channel, time-averaged within each time bin. These calculations yielded 384 neural features per participant, which were used for offline analysis without spike sorting (Trautmann et al., 2019). All features were normalized by subtracting the block-specific mean activity of the features within each recording block, in order to minimize non-stationarities in the data.

Unless otherwise stated, all subsequent offline analyses of neural data were performed using MATLAB software.

### Characterization of Individual Neural Feature Tuning

The first goal of this study was to determine the degree to which force- and grasp-related information are represented within individual TC and SBP neural features. Specifically, neural activity resulting from three discrete *forces* and two (Session 1), four (Sessions 2-4, 6-7), or five (Session 5) *grasps*, resulted in 6, 12 or 15 conditions of interest per session. See Table 1 for a list of grasps included for each individual research session. To visualize individual feature responses to force and grasp, each feature’s peristimulus time histogram (PSTH) was computed for each of these conditions by averaging the neural activity over go cue-aligned trials. These trials were temporally smoothed with a Gaussian kernel (100-ms standard deviation) to aid in visualization.

To determine how many of these individual features were tuned to force and/or grasp, statistical analyses were implemented in MATLAB and with the WRS2 library in the R programming language (Wilcox, 2017). Briefly, features were pre-processed in MATLAB to compute each feature’s mean go-phase deviation from baseline during each trial. Baseline activity was computed by averaging neural activity across multiple rest trials.

In R, the distribution of go-phase neural deviations was found to be normal (analysis of Q-Q plots and Shapiro-Wilk tests, p < 0.05) but heteroskedastic (Levene’s test, p < 0.05), necessitating a 2-way Welch ANOVA analysis to determine neural tuning to force, grasp, and their interaction (p < 0.05). Features exhibiting an interaction between force and grasp were further separated into individual grasp conditions (closed pinch, open pinch, ring pinch, power, elbow), within which one-way Welch-ANOVA tests were implemented to find interacting features that were tuned to force. All p values were corrected for multiple comparisons using the Benjamini-Hochberg procedure (Benjamini and Hochberg, 1995).

### Neural Population Analysis and Decoding

The second goal of this study was to determine the degree to which force and grasp are represented within – and can be decoded from – the level of the neural population. Here, the neural population was represented using both traditional and demixed principal component analysis (PCA).

#### Visualizing Force Representation with PCA

In order to visualize how consistently forces were represented across different grasps, neural activity collected during Sessions 5 and 7 were graphically represented within a low-dimensional space found using PCA. Notably, during Session 5, participant T8 attempted to produce three discrete forces not only with several grasps, but also with an elbow extension movement. Therefore, two sets of PCA analyses were implemented on the data. The first, which was applied to both sessions, performed PCA on all force and grasp conditions within the session. In the second analysis specific to Session 5 only, PCA was applied solely on power grasping and elbow extension trials in order to elucidate whether forces were represented in a consistent way across the entire upper limb. For both analyses, the PCA algorithm was applied to neural feature activity that was averaged over multiple trials and across the go phase of the task.

The results of each decomposition were plotted in a low-dimensional space defined by the first two principal components. The force axis within this space, given by Equation 1, was estimated by applying multi-class linear discriminant analysis (LDA) (Juric, 2020) to the centered, force-labelled PCA data, and then using the largest LDA eigenvector as the multi-dimensional slope ***m*** of the force axis. Here, ***PC_score_*** is the principal component score or representation of the neural data in PCA space and *f* is the intended force level. A consistent force axis across multiple grasps within PCA space would suggest that forces are represented in an abstract (and thus grasp-independent) manner.

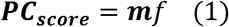

#### Demixed Principal Component Analysis

The remainder of population-level analysis was implemented using demixed principal component analysis (dPCA). dPCA is dimensionality reduction technique that, similarly to traditional PCA, decomposes neural population activity into a few components that capture the majority of variance in the source data (Kobak et al., 2016). Unlike traditional PCA, which yields principal components (PCs) that capture signal variance due to multiple parameters of interest, dPCA performs an ANOVA-like decomposition of data into *task-dependent* dimensions of neural activity. Briefly, the matrix ***X*** of neural data is decomposed into trial-averaged neural activity explained by time (*t*), various task parameters (*T8, T5*), their interaction (*T8T5*), and noise, according to Equation 2. Next, dPCA finds separate decoder (***D***) and encoder (***E***) matrices for each marginalization *M* by minimizing the loss function *L* exhibited in Equation 3.

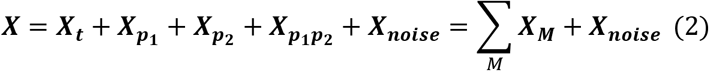

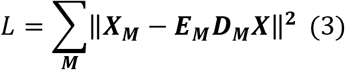

The resulting demixed principal components (dPCs), obtained by multiplying the neural data ***X*** by the rows of each decoder matrix ***D_M_***, are, in theory, de-mixed, in that the variance explained by each component is due to a single, specific task parameter *M*. These dimensions of neural activity not only reveal population-level trends in neural data, but they can also be used to decode task parameters of interest (Kobak et al., 2016).

#### Single dPCA Component Implementation

In the present study, the task parameters of interest were force and grasp. Here, one goal was to use variance as a metric to quantify the degree to which force and grasp were represented within the neural population as a whole. Therefore, for each research session listed in Table 1, the neural data ***X*** was temporally smoothed using a Gaussian filter (100 millisecond standard deviation) and decomposed into neural activity that varied with four marginalizations ***X_M_***, as per Equation 2: *time* (condition independent), *force, grasp*, and an *interaction* between force and grasp. The variance that each marginalization accounted for was computed as the sum of squares of the mean-centered neural data contained within the marginalization.

An additional goal was to isolate neural components that contained useful information about force and grasp, *i.e.,* components that would enable discrimination between individual force levels and grasp types. First, dPCA was used to reduce each of the four, 384-dimensional, mean-centered marginalizations ***X_M_*** into 20 dPCs, as described by Equation 3. This yielded 80 dPCs across all four marginalizations. Second, the variances accounted for by each of the 80 components were computed as the sum of squares. Third, the top 20 out of 80 components with the highest variance were selected as representing the majority of variance in the neural dataset and were assembled into a decoder matrix ***D***. Finally, each of these top 20 components was assigned to one of the four marginalizations of interest according to the marginalization from which it was extracted. For example, dPCs that were extracted from the force marginalization ***X_force_*** were deemed as force-tuned dPCs; those extracted from the grasp marginalization ***X_grasp_*** were deemed as grasp tuned dPCs; and those extracted from the marginalization ***X_F/G_*** representing an interaction between force and grasp were deemed as interacting dPCs.

Each dPC’s information content was further quantified in two ways. First, in order to assess the degree to which dPCs were demixed, each dPC’s variance was subdivided into four sources of variance corresponding to each of the four marginalizations of interest, as per Equation 2. Second, the decoder axis associated with each dPC was used as a linear classifier to decode intended parameters of interest. Specifically, each force-tuned dPC was used to decode force at every time point of the behavioral task, while each grasp-tuned dPC was used to decode grasp, but not force. Likewise, components that exhibited an interaction between force and grasp were used to decode force-grasp pairs. Condition-independent dPCs, which were tuned to time, were not used to decode force or grasp from the neural activity.

Linear classification was implemented using 100 iterations of stratified Monte Carlo leave-group-out cross-validation (Kobak et al., 2016). During each iteration, one random group of *F × G* test “pseudo-trials,” each corresponding to one of the several force-grasp conditions, was set aside during each time point (*F* = number of intended forces, *G* = number of intended grasps). Next, dPCA was implemented on the remaining trials, and the decoder axes of the resulting dPCs were used to predict the intended forces or intended grasps indicated by the test set of pseudo-trials at each time point. This was accomplished by first computing mean dPC values for each force-grasp condition, separately for each time point; projecting the *F × G* “pseudo-trials” onto the decoder axes of the dPCs at each time point; and then classifying the pseudo-trials according to the closest class mean (Kobak et al., 2016). The proportion of *F* × *G* pseudo-trials correctly classified across 100 iterations at each time point constituted a time-dependent classification accuracy. Chance performance was computed by performing 100 shuffles of all available trials, randomly assigning force or grasp conditions to the shuffled data, and then performing the same cross-validated classification procedure within each of the 100 shuffles. Classification accuracies that exceeded the upper range of chance performance were deemed significant.

#### Force and Grasp Decoding Using Multiple dPCs

Two additional goals of this study were to determine whether intended forces could be accurately predicted from neural population data and whether these predictions depended on hand grasp configuration. To this end, dPCs that were tuned to force, grasp, and an interaction between force and grasp were used to construct multi-dimensional force and grasp decoders within each session. Specifically, the force decoder was constructed by combining the decoding axes of force-tuned and interacting components into a single, multi-dimensional decoder ***D_F_***; likewise, the grasp decoder ***D_G_*** was constructed by combining the decoding axes of grasp-tuned and interacting components.

Each of these decoders was used to perform 40 runs of linear force and grasp classification for each of S research sessions per participant, implemented using the aforementioned stratified Monte Carlo leave-group-out cross-validation procedure (S = 6 for T8; S = 1 for T5). As in the single component implementation (Kobak et al., 2016), each run was accomplished in multiple steps. First, the mean values of all dPCs included within the multi-dimensional decoder were computed for each force-grasp condition, separately for each time point. Second, at each time point, the *F × G* “pseudo-trials” were projected onto the multi-dimensional decoder axis and classified according to the closest class mean. The proportion of test trials correctly classified at each time point over 100 iterations constituted a time-dependent force or grasp classification accuracy.

The aforementioned computations yielded 40 × S time-dependent force and grasp classification accuracies per participant. Session-averaged, time-dependent force and grasp classification accuracies were computed by averaging the performance over 240 session-runs for participant T8 (40 runs × 6 sessions) and 40 session-runs for participant T5 (40 runs × 1 session). These averages were compared to chance performance, which was computed by performing 100 shuffles of all trials, randomly assigning force or grasp conditions to the shuffled data, and then performing force and grasp classification on each of the shuffled datasets using the multidimensional decoders ***D_F_*** and ***D_G_***. Time points in which force or grasp classification exceeded the upper bound of chance were deemed to contain significant force-related or grasp-related information.

To visualize the degree to which individual forces and grasps could be discriminated, confusion matrices were computed over go-phase time windows during which the neural population contained significant force- and grasp-related information. The time window began when session-averaged, time-dependent classification accuracy exceeded 90% of maximum achieved performance within the go phase, and ended at the end of the go phase. First, classification accuracies for each of the S × 40 session-runs were approximated by averaging classification performance across the pre-specified go-phase time window. These time-averaged accuracies, which are henceforth referred to as mean force and grasp accuracies, were next averaged over all S × 40 session-runs to yield confusion matrix data. In this way, confusion matrices were computed to visualize force-related discriminability across all trials, force-related discriminability within individual grasp types, and grasp-related discriminability across all trials.

Classification performances for individual forces and individual grasps were statistically compared using parametric tests implemented on mean force and grasp accuracies. Specifically, for each participant, mean classification accuracies for light, medium, and hard forces were compared by implementing one-way ANOVA across mean force accuracies from all S × 40 session runs. The resulting p values were corrected for multiple comparisons using the Benjamini-Hochberg procedure (Benjamini and Hochberg, 1995). Likewise, mean classification accuracies for closed pinch, open pinch, ring pinch, power, and elbow “grasps” were compared by implementing one-way ANOVA across all mean grasp accuracies. These comparisons were implemented to determine whether offline force and grasp decoding yielded similar versus different classification results across multiple forces and multiple grasps.

Statistical analysis was also used to determine the degree to which grasp affected force decoding accuracy. This was achieved by implementing two-way ANOVA on mean force accuracies that were labelled with the grasps that were used to emulate these forces. The results of the two-way ANOVA showed a statistically significant interaction between force and grasp. Therefore, the presence of simple main effects was assessed within each force level and within each grasp type. Specifically, one-way ANOVA was implemented on mean accuracies within individual force levels to determine whether light forces, for example, were classified with similar degrees of accuracies across all grasp types. Similarly, one-way ANOVA was implemented on mean accuracies within individual grasps to see whether intended forces affected grasp classification accuracy. P values resulting from these analyses were corrected for multiple comparisons using the Benjamini-Hochberg procedure.

Finally, this study evaluated how well dPCA force decoders could generalize to novel grasp datasets in T8 Session 5 and T5 Session 7. Specifically, within each session, a multi-dimensional force decoder ***D_F_*** was trained on neural data generated during all but one grasp type, and then its performance was evaluated on the attempted forces emulated using the left-out “novel” grasp. To establish the generalizability of force decoding performance across many novel grasps, this analysis cycled through all available grasps attempted during Session 5 (closed pinch, open pinch, ring pinch, power, elbow extension) and Session 7 (closed pinch, open pinch, ring pinch, power). For each novel grasp, the trained decoder ***D_F_*** was used to perform 40 runs of stratified Monte Carlo leave-group-out cross-validated linear force classification on two sets of test data: the “initial grasp” dataset, which originated from the grasps on which the force decoder was trained; the “novel grasp” dataset, which originated from the leave-out test grasp. The resulting time-dependent, “initial grasp” and “novel grasp” decoding performances from the go-phase time window during above-90% classification accuracy were averaged over 40 runs and then compared using a standard T test. P values resulting from the statistical analysis were corrected for multiple comparisons across forces and test grasps using the Benjamini-Hochberg procedure.

## Results

### Characterization of Individual Neural Features

Figure 2 shows the activity of four exemplary features from session 5 chosen to illustrate tuning to *force*, *grasp*, *both* force and grasp independently, and an *interaction* between force and grasp, as evaluated with 2-way Welch-ANOVA (corrected p < 0.05, Benjamini-Hochberg procedure). These features demonstrate neural modulation to forces that T8 attempted to produce using all five grasp conditions: closed pinch, open pinch, ring pinch, power grasp, and elbow extension. Figure 2–1 shows the activity of four additional features from participant T5. TC features are labelled from 1-192 according to the recording electrodes from which they were extracted. Corresponding SBP features are labelled from 193-384.

**Figure 2.**
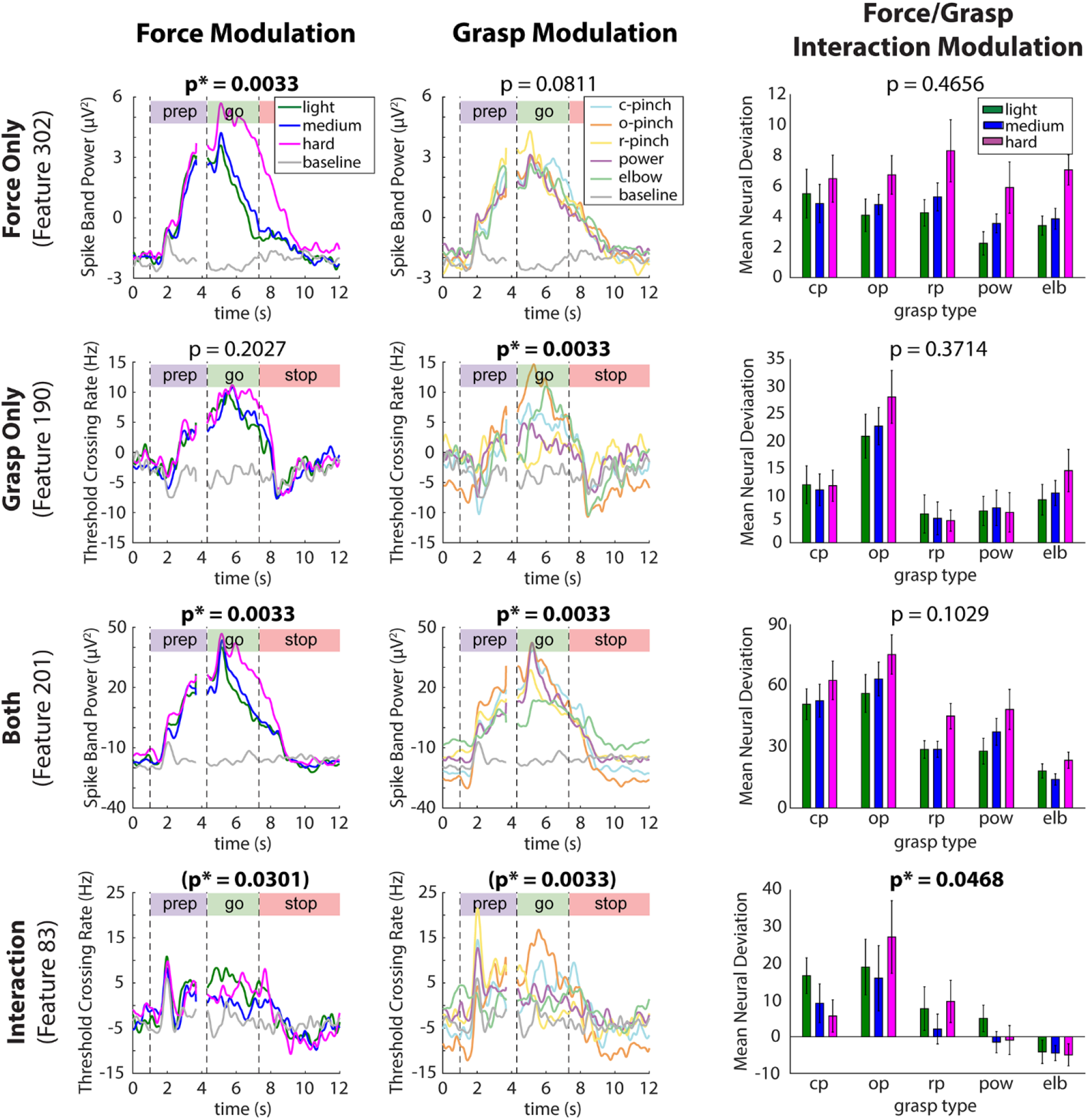
Exemplary threshold crossing (TC) and spike band power (SBP) features tuned to task parameters of interest in participant T8. (TC and SBP features in participant T5 are illustrated in Figure 2–1.) Rows indicate average per-condition activity (PSTH) of four exemplary features tuned to *force*, *grasp*, *both* factors, and an *interaction* between force and grasp, recorded during session 5 from participant T8 (2-way Welch-ANOVA, corrected p < 0.05, Benjamini-Hochberg method). Neural activity was normalized by subtracting block-specific mean feature activity within each recording block, and then smoothed with a 100-millisecond Gaussian kernel to aid in visualization. Column 1 contains PSTHs averaged within individual force levels (across multiple grasps), such that observable differences between data traces are due to force alone. Similarly, column 2 shows PSTHs averaged within individual grasps (across multiple forces). Column 3 shows a graphical representation of the simple main effects as normalized mean neural deviations from baseline activity during force trials within each of the five grasps. Mean neural deviations were computed over the go phase of each trial and subsequently averaged within each force-grasp pair. Error bars indicate 95% confidence intervals.

For each feature, column 1 shows neural activity that was averaged across grasp types (within force levels), resulting in trial-averaged feature traces whose differences in modulation were due to force alone. Similarly, Column 2 shows neural activity averaged within individual hand grasps. Here, SBP feature 302 exhibits modulation to *force only* (row 1), as indicated by statistically significant go-phase differentiation in activity across multiple force levels, but not across multiple grasp levels. This force-only tuning is what might be expected for a “high-level” coding of force that is independent of grasp type. In contrast, TC feature 190 is statistically tuned to *grasp only*, in that it exhibits go-phase differentiation across multiple grasps, but not across multiple forces. SBP feature 201, in which multiple forces and multiple grasps are statistically discriminable, is tuned to *both* force and grasp.

Column 3 of Figure 2 displays a graphical representation of the simple main effects of the 2-way Welch-ANOVA analysis, as shown by mean go-phase neural deviations from baseline feature activity during the production of each individual force level using each individual grasp type. Here, SBP features 302 and 201, which were both tuned to force independent of grasp, showed similar patterns in modulation to light, medium, and hard forces within individual grasp types. In contrast, TC feature 83 was tuned to an *interaction* between force and grasp; accordingly, its modulation to light, medium, and hard forces varied according which grasp type the participant used to emulate these forces. This type of interaction is what might be expected for a more “motoric” encoding of force and grasp type. If each grasp requires a different set of muscles and joints to be active, then a motoric encoding of joint or muscle motion would end up representing force differently depending on the grasp.

Figure 3 summarizes the tuning properties of all 384 TC and SBP neural features in participants T8 and T5, as evaluated with robust 2-way Welch-ANOVA. Specifically, Figure 3A shows the fraction of neural features tuned to *force*, *grasp*, *both* force and grasp, and an *interaction* between force and grasp. Features belonging to the former three groups (i.e., those that exhibited no interactions between force and grasp tuning) were deemed as *independently tuned* to force and/or grasp. As shown in row 1, the proportion of features belonging to each of these groups varied considerably across experimental sessions. However, during all sessions in both participants, a substantial proportion of features (ranging from 15.4-54.7% of the feature population across sessions) were tuned to force, independent of grasp. In other words, a substantial portion of the measured neural population represented force and grasp independently.

**Figure 3.**
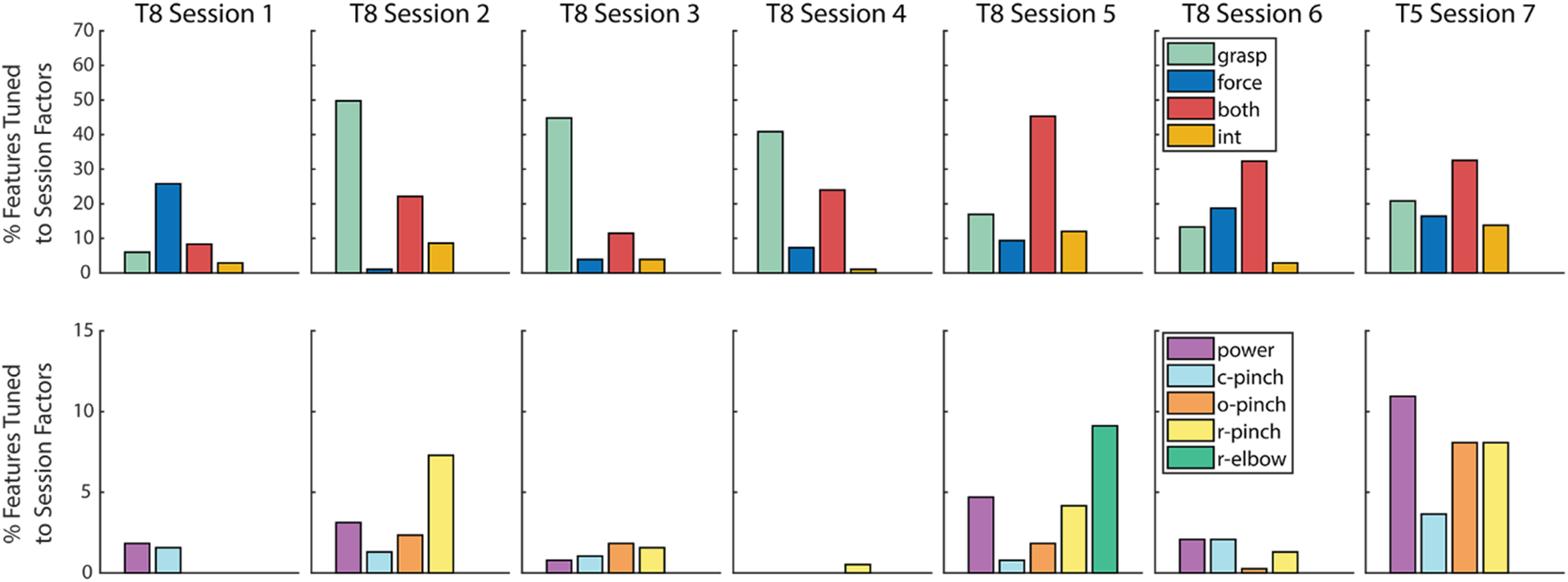
Summary of neural feature population tuning to force and grasp. Row 1: fraction of neural features significantly tuned to *force*, *grasp*, *both* force and grasp, and an *interaction* between force and grasp in participants T8 and T5 (2-way Welch-ANOVA, corrected p < 0.05). Row 2: Fraction of neural features significantly tuned to an interaction between force and grasp, subdivided into force-tuned features within each individual grasp (c-pinch = closed pinch, o-pinch = open pinch, r-pinch = ring pinch). Note that the number of grasp types differed between sessions (see Table 1).

A smaller subset of features exhibited an interaction between force and grasp in both T8 (5.2 +/− 4.2%) and T5 (13.8%). Row 2 of Figure 3 further separates these interacting features into those that exhibited force tuning within each individual grasp type, as evaluated by one-way Welch-ANOVA (corrected p < 0.05). Here, the proportion of interacting features tuned to force appeared to depend on grasp type, particularly during sessions 2, 4, 5, 6, and 7, in a session-specific manner. In other words, within a small contingent of the neural feature population, force representation showed some dependence on intended grasp. Taken together, Figure 3 suggests that force and grasp are represented both independently and dependently within motor cortex at the level of individual neural features.

### Neural Population Analysis and Decoding

#### Simulated Force Encoding Models

The goal of this study was to clarify the degree to which hand grasps affect neural force representation and decoding performance, in light of conflicting evidence of grasp-independent (Chib et al., 2009; Hendrix et al., 2009; Pistohl et al., 2012; Intveld et al., 2018) versus grasp-dependent (Hepp-Reymond et al., 1999; Degenhart et al., 2011) force representation in the literature. Prior to visualizing population-level representation of force, we first illustrate these differing hypotheses with a toy example of expected grasp-independent versus grasp-dependent (interacting) representations of force within the neural space. Figure 4 simulates grasp-independent force encoding with an additive model, given by Equation 4, and grasp-dependent force encoding with a scalar model, given by Equation 5.

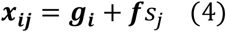

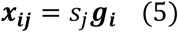

**Figure 4.**
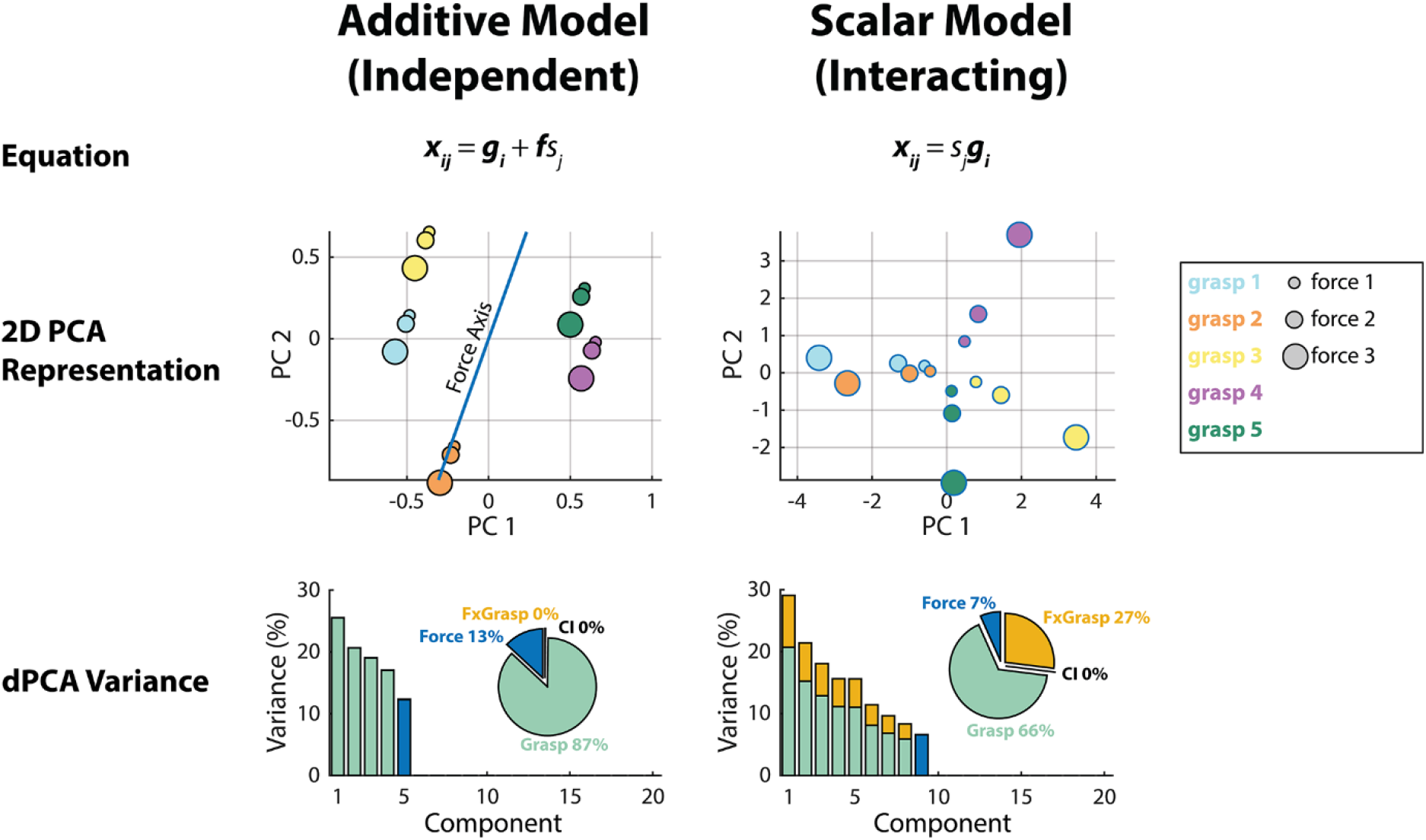
Simulated models of independent and interacting (grasp-dependent) neural representations of force. Row 1: Equations corresponding to the independent and interacting models of force representation. Here, ***x_ij_*** represents neural feature activity generated during a particular grasp *i* and force *j*, ***g_i_*** represents baseline feature activity during grasp *i*, ***f*** represents force-related neural feature activity, and *sj* is a discrete force level. Row 2: Simulated population neural activity projected into a two-dimensional PCA space. Estimated force axes within the low-dimensional spaces are shown as blue lines. Row 3: Summary of variances accounted for by the top 20 demixed principal components extracted from the simulated neural data from each model. Here, the variance of each individual component is separated by marginalization (*force, grasp,* and *interaction* between force and grasp). Pie charts indicate the percentage of total signal variance due to these marginalizations.

In these equations, ***x_ij_*** is a vector of trial-averaged activity from 100 simulated neural features, generated during a particular grasp *i* and force *j*. Here, ***g_i_*** is a 100 × 1 vector of normalized baseline feature activity during the grasp *i*, ***f*** is a 100 × 1 vector of normalized baseline neural feature activity during force generation, and *s_j_* is a discrete, scalar force level (1, 2 or 5). The vectors ***g_i_*** and ***f*** contained values drawn from the standard normal distribution.

Within the additive model in Equation 4, the overall neural activity ***x_ij_*** is represented as a summation of independent force- and grasp-related contributions. Thus, Equation 4 models independent neural force representation, in which force is represented at a high level independent of grasp. In contrast, the scalar encoding model in Equation 5 models the neural activity ***x_ij_*** as resulting from a multiplication of the force level *s_j_* and the baseline grasping activity ***g_i_***. Such an effect might be expected if force were encoded as low-level tuning to muscle activity. In this case, different force levels would result in the same pattern of muscle activity being activated to a lesser or greater degree, thus scaling the neural activity associated with that grasp, resulting in a coupling between force and grasp. Therefore, Equation 5 models an interacting (grasp-dependent) neural force representation.

Row 2 of Figure 4 shows simulated neural activity resulting from the independent and interacting encoding models within two-dimensional PCA space. In the independent model, force is represented in a consistent way across multiple simulated grasps, as indicated by the force axis. In contrast, within the interacting model, force representation differs according grasp. These differences are further highlighted in Row 3 of Figure 4, in which dPCA was applied to the simulated neural data (over 20 simulated trials) resulting from each model. While the additive model exhibited no interaction-related neural variance, the scalar model yielded a substantial proportion of force, grasp, and interaction-related variance. Note that within these toy models, the simulated neural activity did not vary over its time course and, thus, exhibited no condition-independent (time-related) variance.

#### Neural Population Analysis

Figure 5 shows neural population-level activity patterns during two sessions from participants T8 and T5. In the first two columns, dPCA and traditional PCA were applied to all force-grasp conditions in both participants. In the third column, these dimensionality reduction techniques were applied solely to force trials attempted using the power grasping and elbow extension, in order to further quantify force representation across the entire upper limb.

**Figure 5.**
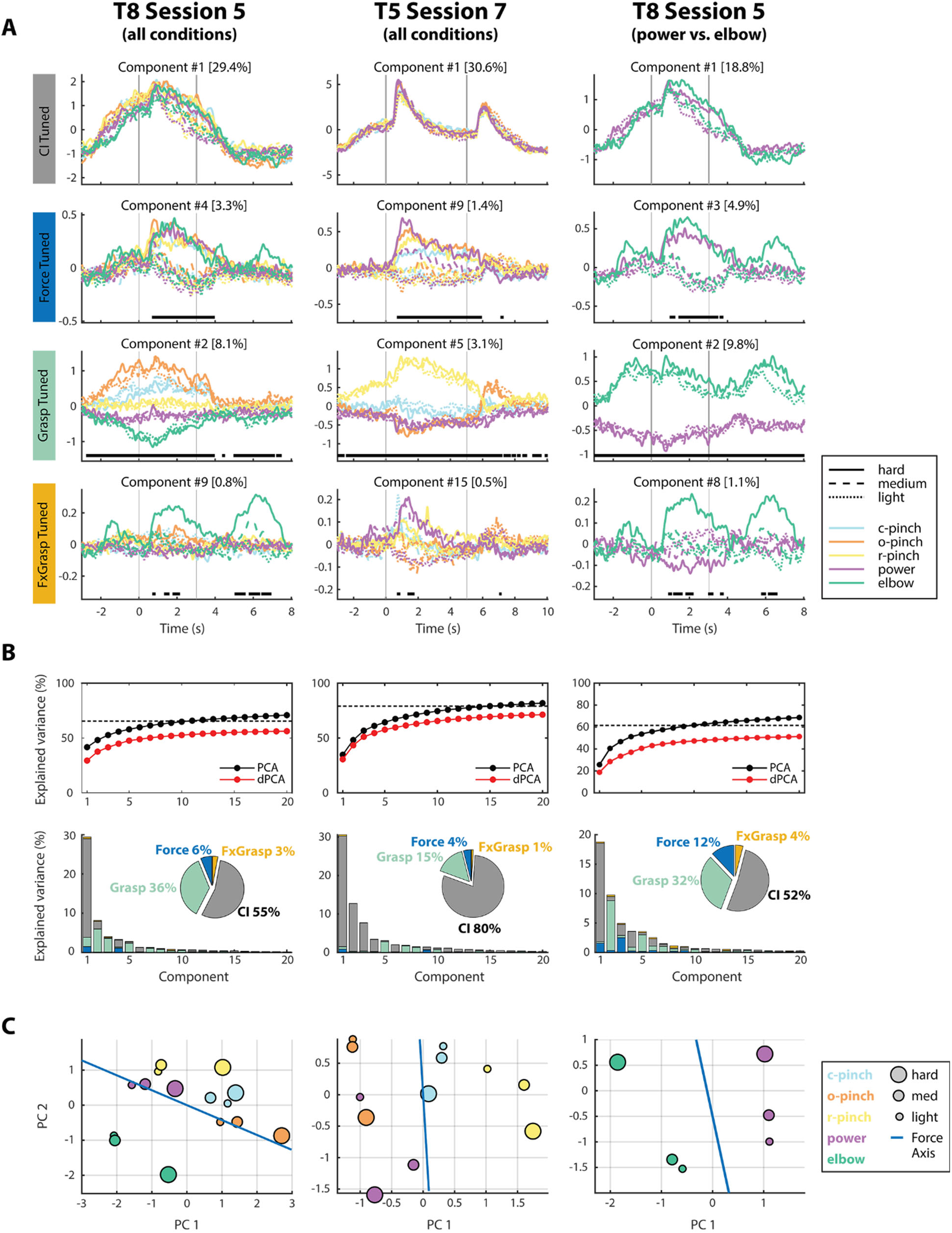
Neural population-level activity patterns. **A.** Demixed principal components (dPCs) isolated from all force-grasp conditions from T8 Session 5, all force-grasp conditions from T5 Session 7, and power versus elbow conditions from T8 Session 5 neural data. (Additional sessions are shown in Figure 5–1). For columns 1 and 2, dPCs were tuned to four marginalizations of interest: *time* (condition-independent tuning), *force, grasp*, and an *interaction* between force and grasp (f/grasp tuning). For column 3, dPCs were tuned to *time, force, movement,* and an *interaction* between force and movement (f/movement tuning). dPCs that account for the highest amount of variance in the per-marginalization neural activity are shown here. These variances are included in brackets next to each component number. Vertical bars indicate the start and end of the go phase. Horizontal bars indicate time points at which the decoder axes of the pictured components classified forces (row 2), grasps/movements (row 3), or force-grasp/force-movement pairs (row 4) significantly above chance. **B.** Summary of variances accounted for by the top 20 dPCs and PCs from each session. Here, the variance accounted for by the dPCs approaches the variance accounted for by traditional PCs. Horizontal dashed lines indicate total signal variance, excluding noise. Row 2 shows the variance of each individual component, separated by marginalization **C.** Go-phase activity within a two-dimensional PCA space. Estimated force axes within the low-dimensional PCA spaces are shown as blue lines.

The twelve dPCs shown in Figure 5A explain the highest amount of variance within each of the four marginalizations of interest, for each participant. For example, participant T8’s Component #4 (row 2, column 1) is the largest force-tuned component in the dataset and explains 3.3% of the neural data’s overall variance. Similarly, T8’s Component #2 (row 3, column 1), which captures grasp-related activity, explains 8.1% of neural variance. Horizontal black bars on each panel indicate time points at which individual dPC decoding axes predict intended forces (row 2), grasps (row 3), and force-grasp pairs (row 4) more accurately than chance performance. In both participants, single components were able to offline-decode intended forces at above-chance levels solely during the active “go” phase of the trial, indicated by the vertical gray lines. However, grasp-tuned components were able to accurately predict intended grasps at nearly all time points during the trial, including the prep and stop phases. These trends were observed when dPCA was applied across all force-grasp conditions (Columns 1 and 2) and across solely power and elbow trials in participant T8 (Column 3).

Figure 5B summarizes the variance accounted for by the entire set of dPCs extracted from each dataset. Specifically, the first row shows the cumulative variance captured by the dPCs (red), as compared to components extracted with traditional PCA (black). Here, dPCs extracted from different marginalizations were not necessarily orthogonal, and accounted for less cumulative variance than traditional PCs because the axes were optimized for demixing in addition to just capturing maximum variance. However, the cumulative dPC variance approached total signal variance, as indicated by the dashed horizontal lines in each panel, and were thus deemed as a faithful representation of the neural population data.

The second row of Figure 5B further subdivides the variances of individual dPCs into per-marginalization variances. Here, most of the variance in each extracted component can be attributed to one primary marginalization, indicating that the extracted components are fairly well demixed. Pie charts indicate the percentage of total signal variance (excluding noise) due to force, grasp, force/grasp interactions, and condition-independent signal components. In both participants, condition-independent components accounted for the highest amount of neural signal variance, followed by grasp, then force, then force-grasp interactions. In other words, more variance could be attributed to putative grasp representation than force representation at the level of the neural population. Additionally, force-grasp and force-movement interactions only accounted for a small amount of neural variance, even when dPCA was applied solely across power grasping and elbow extension trials (Column 3). Similar results were found when analyzing additional sessions, as shown in Figure 5–1.

Finally, Figure 5C visualizes the trial-averaged, go-phase-averaged neural activity from each dataset within two-dimensional PCA space. Within these plots, each data point represents the average neural activity corresponding to an individual force-grasp condition. In all panels, light, medium, and hard forces, represented as different shapes within PCA space, aligned to a consistent force axis (shown in blue) across multiple grasps – and also across power grasping and elbow extension movements.

The findings exhibited within Figures 5B and 5C closely resemble the simulation results from the additive force encoding model (Figure 4, Equation 4), which would be expected for grasp-independent force representation. However, these results differ slightly from those expected from the abstract model, in that some amount of interaction-related variance was present in Figure 5B, and that the force activity patterns in Figure 5C deviated to a small degree from the force axis. These small deviations somewhat resemble the scalar encoding model (Figure 4, Equation 5), which would be expected for interacting force and grasp representations.

#### Time-Dependent Decoding Performance

Figure 6 summarizes the degree to which intended forces and grasps could be predicted from the neural activity using the aforementioned dPCs. Here, offline force decoding accuracies were computed by using a force decoder ***D_F_*** – created by assembling the decoding axes of multiple force-tuned and interacting components – to classify light, medium, and hard forces over multiple session-runs of a 100-fold, stratified, leave-group-out Monte Carlo cross-validation scheme, as described in the Methods. Similarly, grasp decoding accuracies in row 3 were computed using a grasp decoder ***D_G_***, created by assembling the decoding axes of grasp-tuned and interacting dPCs. Row 1 of Figure 6 shows time-dependent force decoding results, averaged over S × 40 session-runs in participants T8 (S = 6) and T5 (S = 1). Row 2 further subdivides the results of Row 1 into force decoding accuracies achieved during individual hand grasps. Finally, Row 3, shows time-dependent grasp decoding results for both participants.

**Figure 6.**
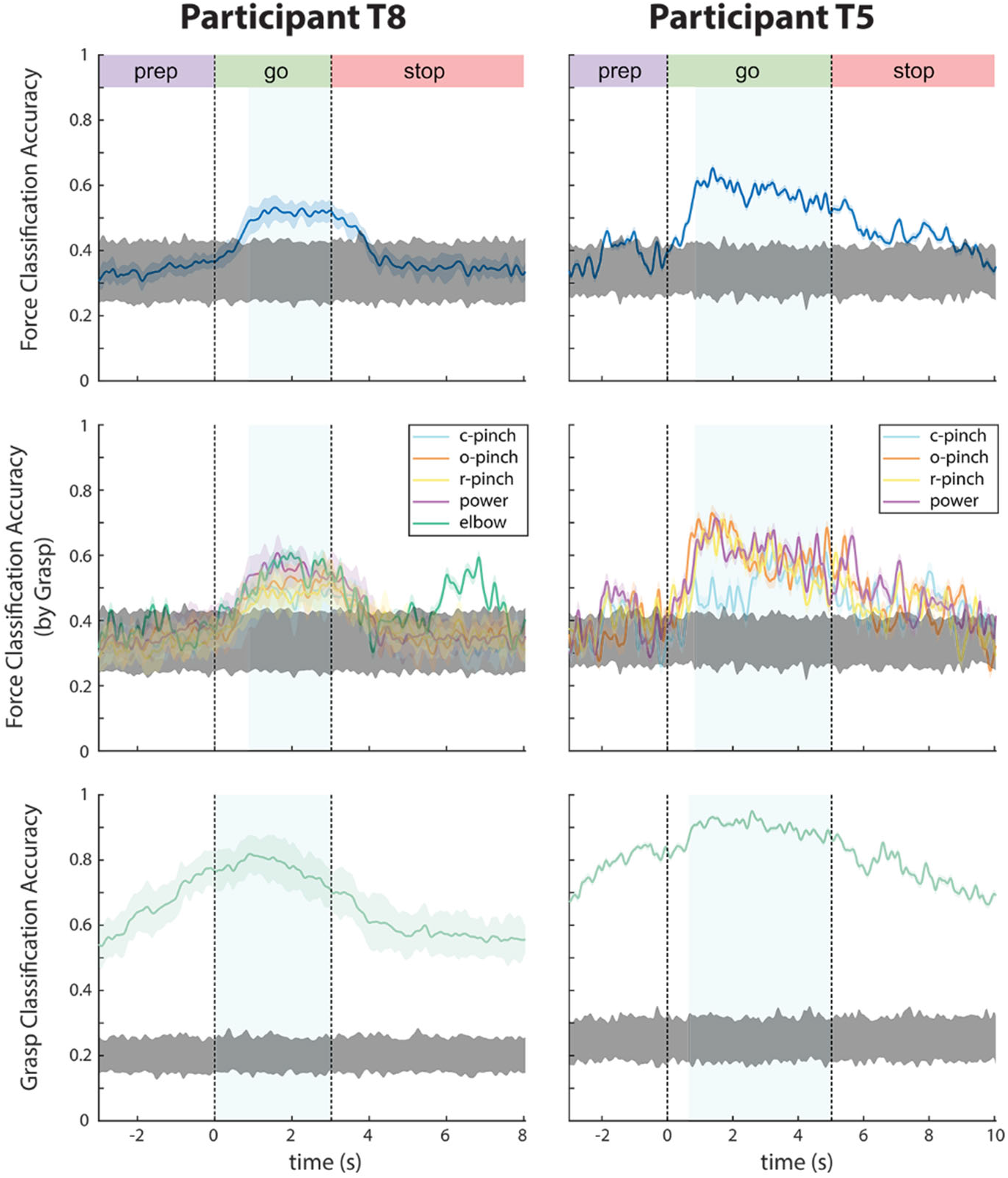
Time-dependent classification accuracies for force (rows 1-2) and grasp (row 3). Data traces were smoothed with a 100 millisecond boxcar filter to aid in visualization. Shaded areas surrounding each data trace indicate the standard deviation across 240 session-runs for most trials in participant T8, 40 session-runs for elbow extension trials in participant T8, and 40 session-runs in participant T5. Gray shaded areas indicate the upper and lower bounds of chance performance over S × 100 shuffles of trial data, where S is the number of sessions per participant. Time points at which force or grasp is decoded above the upper bound of chance are deemed to contain significant force- or grasp-related information. Blue shaded regions indicate the time points used to compute go-phase confusion matrices in Figure 7.

**Figure 7.**
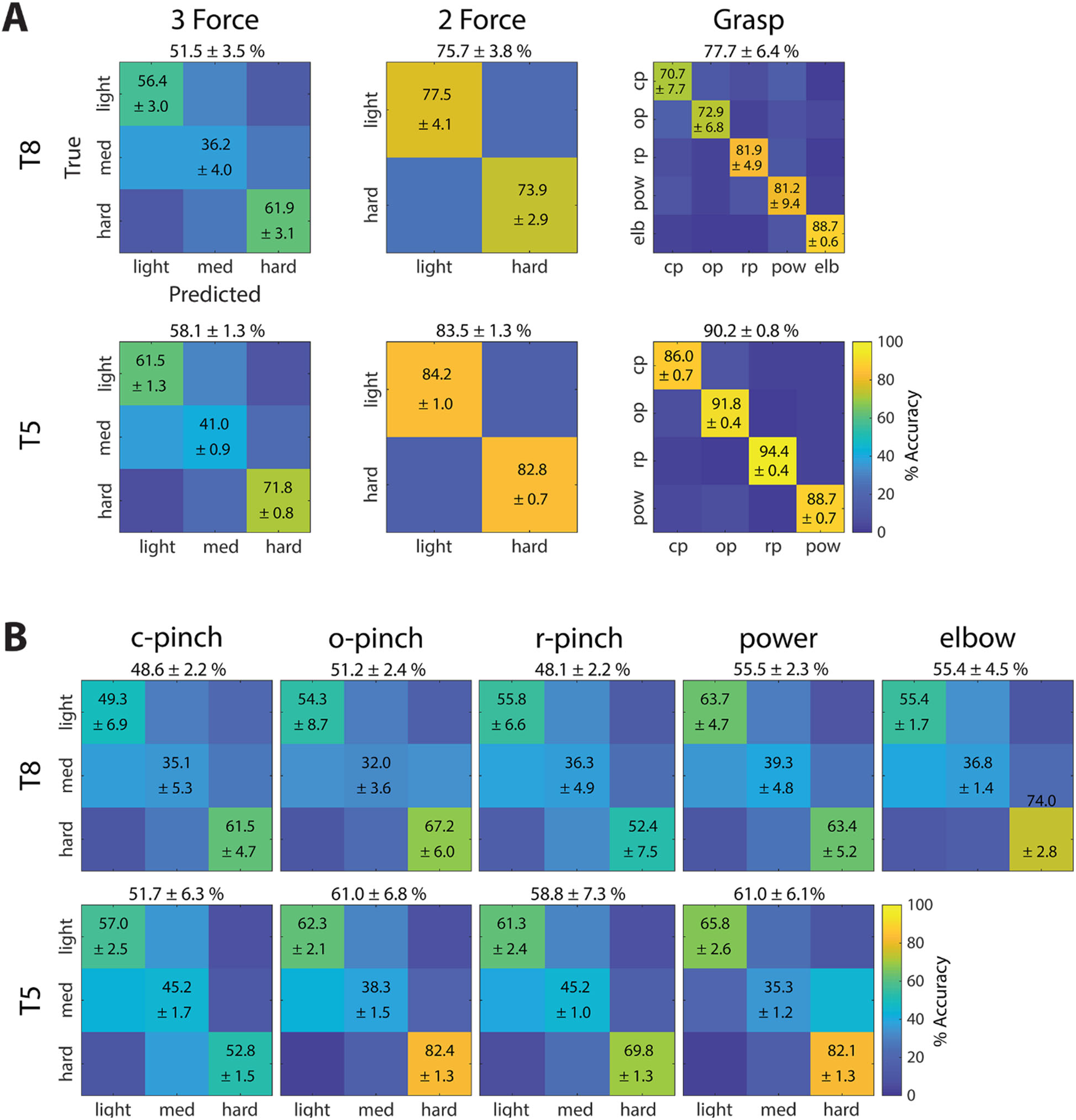
Go-phase confusion matrices. **A.** Time-dependent classification accuracies (shown in Figure 6) were averaged over go-phase time windows that commenced when performance exceeded 90% of maximum, and ended with the end of the go phase. These yielded mean trial accuracies, which were then averaged over all session-runs in each participant. Overall force and grasp classification accuracies are indicated above each confusion matrix. Standard deviations across multiple session-runs are indicated next to mean accuracies (cp = closed pinch, op = open pinch, rp = ring pinch, pow = power, elb = elbow extension). Statistical comparisons between the achieved classification accuracies are shown in Figure 7–1. **B.** Confusion matrices now separated by the grasps that participants T8 (row 1) and T5 (row 2) used to attempt producing forces. Statistical comparisons between the achieved force accuracies are shown in Figures 7–2 and 7–3.

Here, intended forces were decoded at levels exceeding the upper bound of chance solely during the go phase, regardless of the grasp used to emulate the force. The exception to this trend occurred during elbow extension trials, in which intended forces were decoded above chance during the stop phase. In contrast, intended grasps were decoded above chance during all trial phases, regardless of the number of grasps from which the decoder discriminated (Figure 6–2) – though go-phase grasp decoding accuracies tended to exceed those achieved during other trial phases. In summary, both intended forces and grasps were decoded above chance during time periods when participants intended to produce these forces and grasps – and in some cases, during preparatory and stop periods. Time dependent decoding accuracies for individual force levels and individual grasp types are displayed in Figure 6–1.

Time-dependent classification accuracies for individual force levels and grasp types are shown in Figure 6–1. Grasp classification accuracies, separated by number of attempted grasp types, are presented in Figure 6–2.

#### Go-Phase Decoding Performance

Figure 7 summarizes go-phase force and grasp decoding accuracies as confusion matrices. Here, time-dependent classification accuracies for each force level and each grasp type were averaged over go-phase time windows (see Figure 6) that commenced when overall classification performance exceeded 90% of their maximum, and ended with the end of the go phase. This time period was selected in order to exclude the rise time in classification accuracy at the beginning of the go phase, so that the resulting mean trial accuracies reflected stable values. The mean trial accuracies were then averaged over all session-runs in each participant to yield confusion matrices of true versus predicted forces and grasps. Figure 7B further subdivides overall three-force classification accuracies into force classification accuracies achieved during each individual grasp type (columns) in both participants (rows). The confusion matrices in Figure 7 represent cumulative data across multiple sessions in participant T8, and one session in participant T5. Figures 7–1, 7–2 and 7–3 statistically compare decoding accuracies between individual force levels and grasp types within each individual session.

In Figures 6A, 7A, and 6–1, overall three-force classification accuracies exceeded the upper limit of chance in both participants. However, the decoding accuracies of individual force levels were statistically different. For almost all sessions, hard forces were classified more accurately than light forces (with the exception of Session 4, during which light and hard force classification accuracy was statistically similar); and both light and hard forces were always classified more accurately than medium forces. More specifically, hard and light forces were decoded above chance across all sessions, while medium force classification accuracies often failed to exceed chance in both participants.

In contrast, both overall and individual grasp decoding accuracies always exceeded the upper limit of chance. According to Figures 7A and 7–1B, certain grasps were decoded more accurately than others. Specifically, in participant T8, the power and ring pincer grasps were often classified more accurately than the open and closed pincer grasps across multiple sessions (corrected p << 0.05, one-way ANOVA). Elbow extension, which required the participants to attempt force production in the upper limb in addition to the hand, was classified more accurately than any of the grasping forces during Session 5 (corrected p << 0.05). In participant T5, grasp classification accuracies, in order from greatest to least, were ring pincer > open pincer > power > closed pincer. Regardless, grasp decoding performance always exceeded force decoding performance in both participants, as seen in Figures 6 and 7.

In Figures 7 and 7–3, overall and individual force classification accuracies varied depending on the hand grasps used to attempt these forces. Specifically, classification accuracies for forces attempted with different grasps were, with few exceptions, statistically different (corrected p << 0.05, one-way ANOVA). For example, in Figures 7B and 7–3, hard forces attempted using the open pincer grasp were always classified more accurately than hard forces attempted using the ring pincer grasp in both participants. In other words, grasp type affected how accurately forces were decoded.

Finally, Figure 8 summarizes how well force decoders trained on one set of grasps generalize to novel grasp types in T8 Session 5 (row 1) and T5 Session 7 (row 2). A force decoder was used to discriminate forces amongst a set of grasps used for training (“left-in”, gray bars) or a leave-out “novel” grasp (white bars). Here, the force decoding performance between the leave-in and leave-out grasps was significantly different in 7 out of 9 comparisons, suggesting that grasp affects how well forces are decoded from neural activity. However, for all sets of grasps, force decoding performance always exceeded chance. This was even true when, during T8 Session 5, the force decoder was trained on four hand grasps and evaluated on elbow extension data. This is consistent with the previous population-level analyses that show that components of force representation in motor cortex are conserved across grasps and even arm movements.

**Figure 8.**
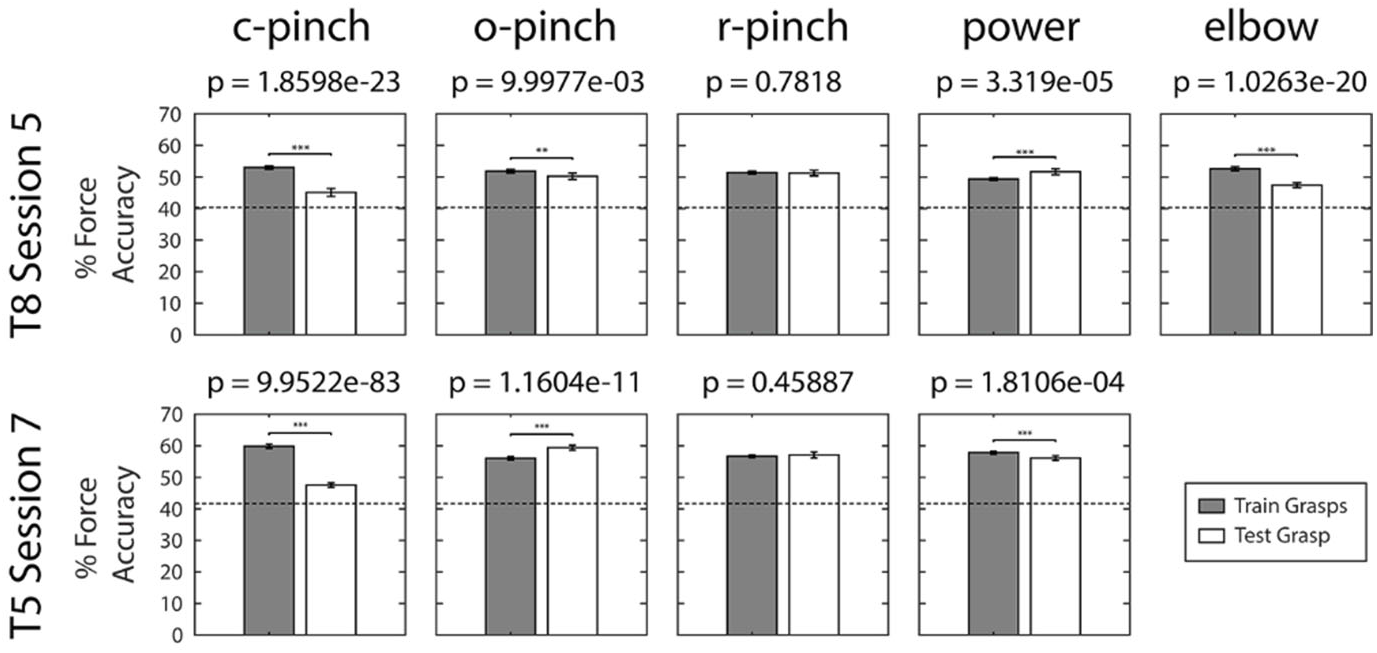
Go-phase force classification accuracy for novel (test) grasps. Within each session (rows), dPCA force decoders were trained on neural data generated during all grasps, excluding a single leave-out grasp type (columns). The force decoder was then evaluated over the set of training grasps (gray bars), as well as the novel leave-out grasp type (white bars). The horizontal dotted line in each panel indicates upper bound of the empirical chance distribution for force classification.

## Discussion

The current study sought to determine how human motor cortex encodes hand grasps and discrete forces, the degree to which these representations overlapped and interacted, and how well forces and grasps could be decoded. Three major findings emerged from this work. First, force information was present in – and could be decoded from – intracortical neural activity in a consistent way across multiple hand grasps. This suggests that force is, to some extent, represented at a high level, independent of motion and grasp. However, as a second finding, grasp affected force representation and classification accuracy. This suggests that there is a simultaneous, low-level, motoric representation of force. Finally, hand grasps were classified more accurately and explained more neural variance than forces. These three findings and their implications for future online force decoding efforts are discussed here.

### Force and Grasp Representation in Motor Cortex

#### Force information persists across multiple hand grasps in individuals with tetraplegia

##### Overall Force Representation

Force was represented in a consistent way across multiple hand grasps within the neural activity. In particular, a substantial contingent of neural features was tuned to force independent of grasp (Figure 3); force-tuned components explained more population-level variance than components tuned to force-grasp interactions (Figure 5); intended forces were accurately predicted from population-level activity across multiple grasps (Figures 6–7); and force decoding performance generalized to novel grasps (Figure 8). The study results suggest that, to a large extent, force is represented at a high level within motor cortex, distinct from grasp, in accordance with the independent force encoding model described by Equation 4 (Figure 4). This conclusion agrees with previous motor control studies (Mason et al., 2004; Chib et al., 2009; Casadio et al., 2015), which suggest that at the macroscopic level, force and motion may be represented independently. In particular, Chib and colleagues showed that descending commands pertaining to force and motion could be independently disrupted via TMS, and that these commands obeyed simple linear superposition laws when force and motion tasks were combined.

Furthermore, intracortical non-human primate studies (Mason et al., 2006; Hendrix et al., 2009; Intveld et al., 2018) suggest that forces are encoded largely independently of the grasps used to produce them. However, in those studies and within the present work, the hand grasps used to produce forces likely recruited overlapping sets of muscle activations. Thus, the relatively low degree of interactions observed here and in the literature could actually be due to overlapping muscle activations rather than truly grasp-independent force representation. For this reason, participant T8 emulated forces using elbow extension in addition to the other hand grasps during Session 5. In Column 3 of Figure 5, dPCA was implemented solely on force trials emulated using elbow extension and power grasping, which involved sets of muscles that operated relatively independently. The resulting dPCA composition yielded a slightly larger amount of variance due to interaction (4%) that was nonetheless smaller than that attributed to force (~12%) or grasp (~35%). Furthermore, discrete force data, when represented within two-dimensional PCA space, aligned closely with a force axis that was conserved over both power grasping and elbow extension movements. These data provide further evidence that force may be encoded independently of movements and grasps.

##### Representation of Discrete Forces

While overall force accuracies exceeded chance performance (Figure 6), hard and light forces were classified more accurately than medium forces across all hand grasps, sessions and participants. In fact, medium forces often failed to exceed chance classification performance (Figures 7A, 6–1B).

Notably, classification performance likely depended on the participants’ ability to kinesthetically attempt various force levels and grasps without feedback, despite having tetraplegia for several years prior to study enrollment. Anecdotally, participant T8 reported that light and hard forces were easier to attempt than medium forces, because they fell at the extremes of the force spectrum and could thus be reproduced consistently. Though his confidence with reproducing all forces improved with training, it is conceivable that, without sensory feedback, medium forces were simply more difficult to emulate cognitively, and thus yielded neural activity patterns that were more inconsistent and difficult to discriminate.

Additionally, this and prior studies suggest that neural activity increases monotonically with increasing force magnitude (Evarts, 1969; Thach, 1978; Cheney and Fetz, 1980; Wannier et al., 1991; Ashe, 1997; Cramer et al., 2002). As a result, medium forces, by virtue of being intermediate to light and hard forces, may be represented intermediate to light and hard forces in the neural space, and may thus be more easily confused with other forces during classification. In the present work, population-level activity associated with medium and light forces appeared similar (Figures 5, 5–1). Likewise, medium forces were most often confused with light forces during offline force classification (Figure 7). The decoding results are consistent with previous studies (Murphy et al., 2016; Downey et al., 2018), in which intermediate force levels were more difficult to discriminate than forces at the extremes of the range evaluated.

#### Hand posture affects force representation and force classification accuracy

##### Single-Feature Versus Population Interactions between Force and Grasp

As previously stated, force information was neurally represented, and could be decoded, across multiple hand grasps (Figures 3, 5–8). However, hand grasp also appears to influence how force information is represented within and decoded from motor cortex. For example, grasp affected how accurately light, medium, and hard forces were predicted from neural activity (Figures 7B, 7–3). Furthermore, despite small force-grasp interaction population-level variance (Figures 5B, 5–1B), as many as 12.0% and 13.8% of neural features exhibited tuning to these interaction effects in participants T8 and T5, respectively (Figure 3), providing further evidence that the force and grasp representation are not entirely independent.

When considering the relatively large number of interacting features and the small population-level interaction variance, one might initially conclude that a discrepancy exists between feature- and population-level representation of forces and grasps. However, we note that the amount of variance explained by a parameter of interest does not always correspond directly to the percentage of features tuned to this parameter. Here, the interaction effects detected within individual features likely reached statistical significance with small effect size. In other words, while real interaction effects were present, as shown in the feature data (Figure 3), the overall effect was small, as exhibited within the population activity (Figure 5). From this perspective, the seemingly incongruous feature- and population-level results actually complement one another and inform our understanding of how forces are represented in motor cortex.

#### Force and Grasp Have Both Abstract (Independent) and Motoric (Interacting) Representations in Cortex

Thus far, studies of force versus grasp representation have largely fallen into two opposing groups. The first proposes that motor parameters are represented independently (Carmena et al., 2003; Mason et al., 2006; Hendrix et al., 2009; Intveld et al., 2018). Such representation implies that the motor cortex encodes an action separately from its intensity, then combines these two events downstream in order to compute the EMG patterns necessary to realize actions in physical space.

In contrast, the second group suggests that force, grasp, and other motor parameters interact within the neural space (Hepp-Reymond et al., 1999; Degenhart et al., 2011). They propose that motor parameters cannot be fully de-coupled (Kalaska, 2009; Branco et al., 2019), and that it may be more effective to utilize the entire motor output to develop a comprehensive mechanical model, rather than trying to extract single parameters such as force and grasp (Ebner et al., 2009).

The current study presents evidence supporting both independent and interacting representations of force and grasp. These seemingly contradictory results actually agree with a previous non-human primate study that recorded from motor areas during six combinations of forces and grasps (Intveld et al., 2018). Intveld and colleagues found that, while force-grasp interactions explained only 0-3% of population variance, roughly 10-20% of recorded neurons exhibited such interactions. These results are highly consistent those outlined in the present study (Figures 3, 5). Thus, the neural space could consist of two contingents: one that encodes force at a high level independent of grasp and motion, and another that encodes force as low-level tuning to muscle activity, resulting in interactions between force and grasp. The second contingent, however small, significantly impacts how accurately forces and grasps are decoded (Figures 7B, 7–3), and should thus not be discounted.

#### Hand grasp is represented to a greater degree than force at the level of the neural population

##### Go-Phase Grasp Representation

In the present datasets, grasps were decoded more accurately (Figures 6–7, 6–1B) and explained more signal variance (Figures 5B, 5–1B) than forces. This suggests that within the sampled region of motor cortex, grasp is represented to a greater degree than force, which agrees with prior literature (Hendrix et al., 2009; Intveld et al., 2018).

Previous studies suggest several reasons why force may be represented to a lesser degree than grasp in the current work. First, force information may have stronger representation in caudal M1, particularly on the banks of the central sulcus (Kalaska and Hyde, 1985; Sergio et al., 2005; Hendrix et al., 2009). Second, force-tuned neurons in motor cortex respond more to the direction of applied force rather its magnitude (Kalaska and Hyde, 1985; Kalaska et al., 1989; Taira et al., 1996). Finally, intracortical non-human primate studies (Georgopoulos et al., 1983; Georgopoulos et al., 1992) and fMRI studies in humans (Branco et al., 2019) suggest that motor cortical neurons respond more to the dynamics of force than to static force tasks. The present work, which recorded from rostral motor cortex and studied the representation of static, non-directional forces, may therefore have detected weaker force-related representation than would have been possible from more caudally-placed recording arrays during a dynamic, functional force task.

Additionally, both study participants were paralyzed and deafferented and received no sensory feedback regarding the forces and grasps they attempted. Previous work suggests that in individuals with tetraplegia, discrepancies exist between the representation of kinematic parameters such as grasp – which remain relatively intact due to their reliance on visual feedback – and kinetic parameters such as force (Rastogi et al., 2020). Specifically, since force-related representation relies heavily on proprioceptive and tactile feedback (Tan et al., 2014; Tabot et al., 2015; Schiefer et al., 2018), whose neural pathways are altered during tetraplegia (Solstrand Dahlberg et al., 2018), the current study may have yielded weaker force-related representation than if this feedback had been included. Therefore, further investigations of force representation are needed in individuals with tetraplegia during naturalistic, dynamic tasks that incorporate sensory feedback – either from intact sensation or from intracortical microstimulation (Flesher et al., 2016) – in order to determine the full extent of motor cortical force representation and to maximize force decoding performance.

##### Grasp Representation During Prep and Stop Phases

Unlike forces, which were represented primarily during the active “go” phase of the trial, grasps were represented throughout the entire task (Figures 5–6), even during the preparatory and stop phases. The ubiquitous representation of grasp observed here could be partially explained by the behavioral task. As described in the Methods, research sessions consisted of multiple data collection blocks, each of which was assigned to a particular hand grasp, and cycled through three attempted force levels within each block (Figure 1B). Thus, while attempted force varied from trial to trial, attempted hand grasps were constant over each block and known by participants in advance. When individuals have prior knowledge of one task parameter, but not another other, information about the known parameter can appear within the baseline activity (Vargas-Irwin et al., 2018). Therefore, grasp-related information may have been represented within the neural space during non-active phases of the trial, simply by virtue of being known in advance.

Additionally, the placement of the recording arrays could have influenced grasp representation in this study. As described in the Methods, two microelectrode arrays were placed within the “hand knob” of motor cortex in each participant (Yousry et al., 1997). These arrays may have recorded from “visuomotor neurons,” which modulate both to grasp execution and to the presence of graspable objects prior to active grasp (Carpaneto et al., 2011), or from neurons that are involved with motor planning of grasp (Schaffelhofer et al., 2015). These neurons have typically been attributed to area F5, a homologue of premotor cortex in non-human primates. Indeed, recent literature in a human participant indicates that the precentral gyrus, rather than belonging to primary motor cortex, is actually part of premotor cortex (Willett et al., 2020). Thus, then the arrays in this study likely recorded from premotor neurons, which modulate to grasp during both visuomotor planning and grasp execution, as was observed here.

### Implications for Force Decoding

#### Hand Grasp Affects Force Decoding Performance

Our decoding results demonstrate that, in individuals with tetraplegia, forces can be decoded offline from neural activity across multiple hand grasps (Figures 6–8). These results agree with the largely independent force and grasp representation of force within single features (Figure 3) and the neural population (Figure 5). From a functional standpoint, this supports the feasibility of incorporating force control into real-time iBCI applications. On the other hand, grasp affects how accurately discrete forces are predicted from neural data (Figures 7B, 7–3). Therefore, future robust force decoders may need to account for additional motor parameters, including hand grasp, in order to maximize performance.

#### Decoding Motor Parameters with Dynamic Neural Representation

The present study decoded intended forces from population activity at multiple time points, with the hope that force representation and decoding performance would be preserved throughout the go phase of the task. We found that force-related activity at the single feature (Figures 2, 2–1) and population levels (Figures 5, 5–1) exhibited both tonic and dynamic characteristics. That is, when study participants attempted to produce static forces, neural modulation to force varied with time to some degree.

The observed dynamic characteristics are consistent with previous results in humans (Murphy et al., 2016; Downey et al., 2018; Rastogi et al., 2020). In particular, Downey and colleagues found that force decoding during a virtual, open-loop, grasp-and-transport task was above chance during the grasp phase of the task, but no greater than chance during static attempted force production during the transport phase. These results support the idea that motor cortex encodes changes in force, rather than (or in addition to) discrete force levels themselves (Smith et al., 1975; Georgopoulos et al., 1983; Wannier et al., 1991; Georgopoulos et al., 1992; Picard and Smith, 1992; Boudreau and Smith, 2001; Paek et al., 2015).

However, the presence of tonic elements agrees with intracortical studies (Smith et al., 1975; Wannier et al., 1991), which demonstrated both tonic and dynamic neural responses to executed forces; and fMRI studies (Branco et al, 2019), which demonstrated a monotonic relationship between the BOLD response and static force magnitudes. Moreover, despite the presence of dynamic response elements, offline force classification performance remained relatively stable throughout the go phase (Figures 6, 6–1), suggesting that the tonic elements could allow for adequate real-time force decoding using linear techniques alone. This may be especially true when decoding forces during dynamic functional tasks, which have been shown to elicit stronger, more consistent neural responses within motor cortex (Georgopoulos et al., 1983; Georgopoulos et al., 1992; Branco et al., 2019).

Nonetheless, real-time force decoding would likely benefit from an exploration of a wider range of encoding models. For example, the exploration of a force derivative model, and its implementation within an online iBCI decoder, would be of potential utility.

#### Decoding of Discrete Versus Continuous Forces

The present work continues previous efforts to characterize discrete force representation in individuals with paralysis (Cramer et al., 2005; Downey et al., 2018; Rastogi et al., 2020) by accurately classifying these forces across multiple hand grasps – especially when performing light versus hard force classification (Figure 7). This supports the feasibility of enabling discrete (“state”) control of force magnitudes across multiple grasps within iBCI systems, which would allow the end iBCI user to perform functional grasping tasks requiring varied yet precise force outputs. Perhaps because discrete force control alone would enhance iBCI functionality, relatively few studies have attempted to predict forces along a continuous range of magnitudes. Thus far, continuous force control has been achieved in non-human primates (Carmena et al., 2003) and able-bodied humans (Pistohl et al., 2012; Chen et al., 2014; Flint et al., 2014), but not in individuals with tetraplegia. If successfully implemented, continuous force control could restore more nuanced grasping and object interaction capabilities to individuals with motor disabilities.

However, during the present work (Figures 7, 6–1) and additional discrete force studies (Downey et al, 2018; Murphy et al, 2016), intermediate force levels were often confused with their neighbors, and thus more difficult to decode. Therefore, implementing continuous force control may pose challenges in individuals with tetraplegia. Possibly, enhancing force-related representation in these individuals via aforementioned techniques – including the introduction of dynamic force tasks, closed loop sensory feedback, and derivative force encoding models – may boost overall performance to a sufficient degree to enable continuous force decoding capabilities. Regardless, more investigations are needed to determine the extent to which continuous force control is possible in iBCI systems for individuals with tetraplegia.

### Concluding Remarks

This study found that, while force information was neurally represented and could be decoded across multiple hand grasps in a consistent way, grasp type had a significant impact on force classification accuracy. From a neuroscientific standpoint, these results suggest that force has both grasp-independent and grasp-dependent (interacting) representations within motor cortex in individuals with tetraplegia. From a functional standpoint, they imply that in order to incorporate force as a control signal in human iBCIs, closed-loop force decoders should ideally account for interactions between force and other motor parameters to maximize performance.

