## Supplemental Figures for "The neural representation of force across grasp types in motor cortex of humans with tetraplegia"

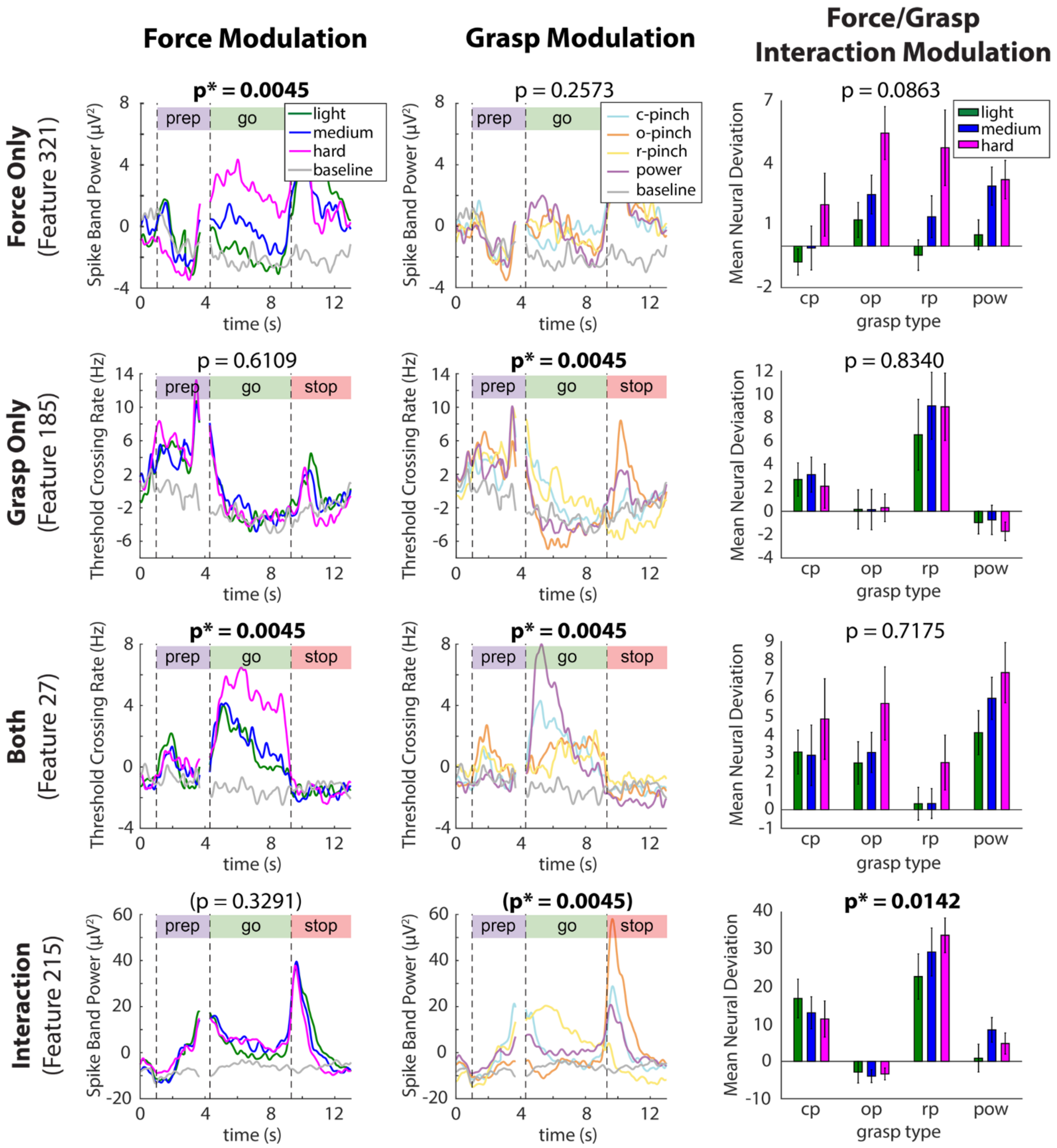


**Figure 2-1.** Exemplary threshold crossing (TC) and spike band power (SBP) features tuned to task parameters of interest in participant T5, presented as in Figure 2. Note the presence of sharp activity peaks during the prep and stop phases of the trial, which were due to the presence of visual cues (Rastogi et al., 2020).

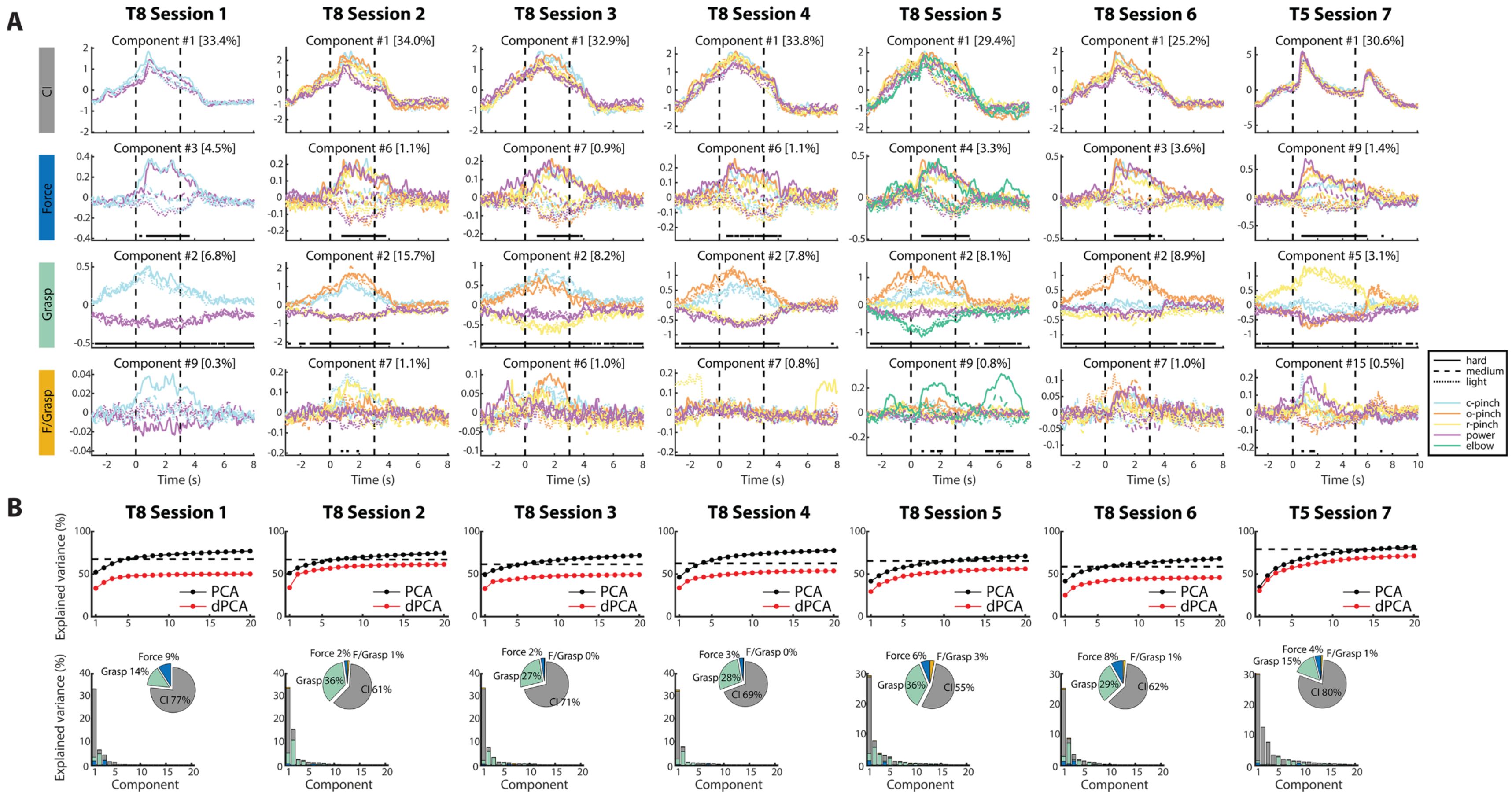


**Figure 5-1.** Neural population-level activity patterns for all sessions, presented as in Figure 5 A-B. **A.** Demixed principal components (dPCs) isolated from all individual sessions of neural data. **B.** Summary of variances accounted for by the top 20 dPCs from each exemplary session.

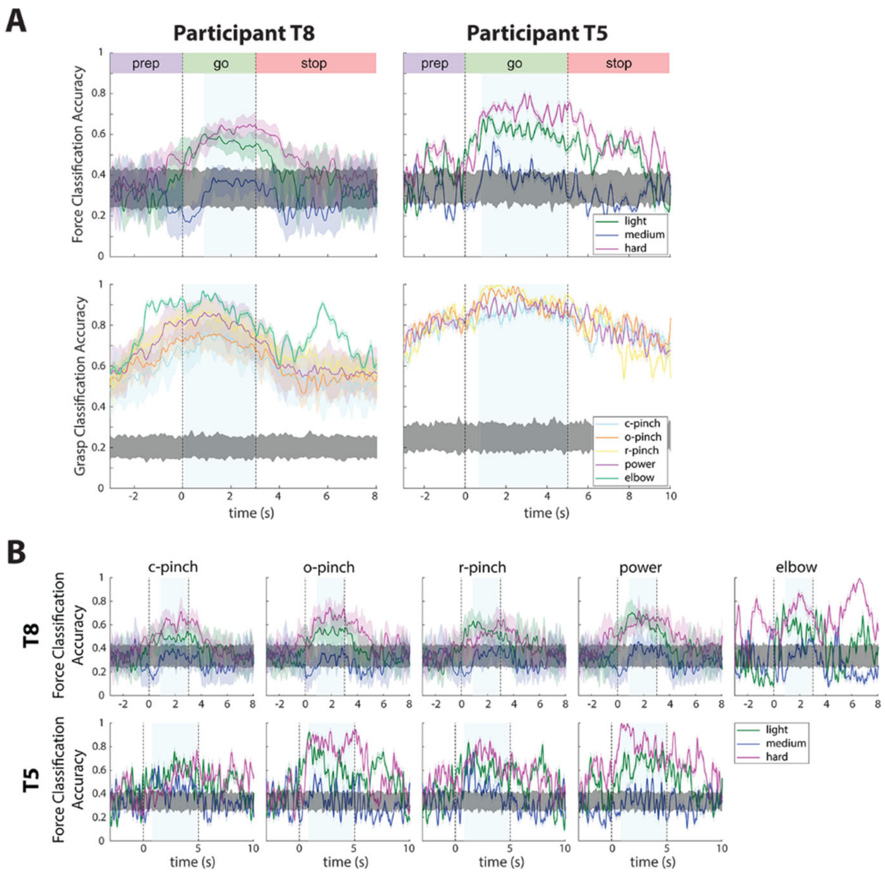


**Figure 6-1.** Time-dependent classification accuracies for individual force levels and grasp types. **A.** Time-dependent classification accuracies for force (row 1) and grasp (row 2), separated by force class and grasp class, respectively. Data traces were smoothed with a 100 millisecond boxcar filter to aid in in visualization. Shaded areas surrounding each data trace indicate the standard deviation across 240 session-runs during most trials in participant T8, 40 session-runs during elbow extension trials in participant T8, and 40-session runs in participant T5. Gray shaded regions indicate the upper and lower bounds of chance performance over S x 100 shuffles of trial data, where S is the number of sessions per participant. Blue shaded regions indicate the time points used to compute go-phase confusion matrices. **B.** Time-dependent force classification accuracies during individual grasps in participants T8 (row 1) and T5 (row 2). Blue shaded regions indicate the time points used to compute go-phase confusion matrices. Decoding performances were averaged over S x 40 session runs, where S is the number of sessions per participant.

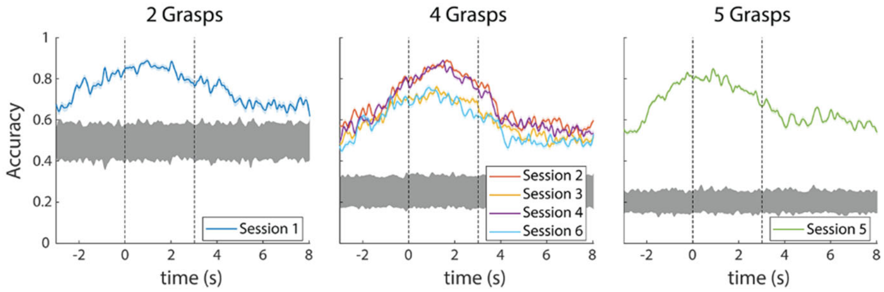


**Figure 6-2.** Time-dependent grasp classification accuracies by number of grasps attempted per session in participant T8. Data traces were smoothed with a 100 millisecond boxcar filter to aid in in visualization. Shaded areas surrounding each data trace indicate the standard deviation across 40 runs during each session in participant T8. Gray shaded regions indicate the upper and lower bounds of chance performance over 100 shuffles of trial data per session. Intended grasp is classified above chance performance at all trial time points, regardless of the number of grasps to be decoded.

A

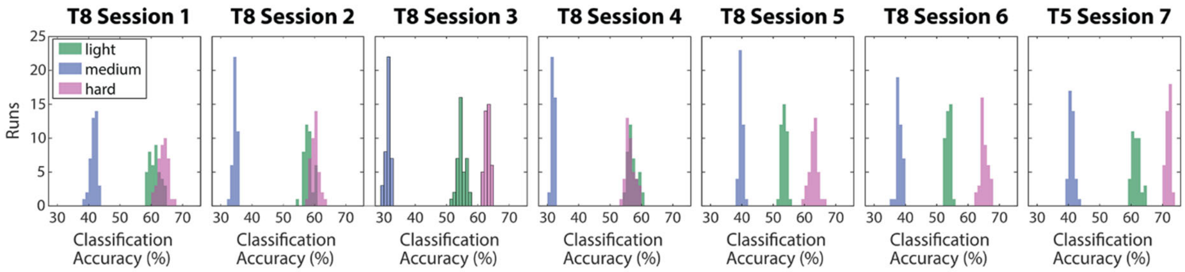


| | $p_1$ | $p_2$ | $p_3$ | $p_4$ | $p_5$ | $p_6$ | $p_7$ |
|---|---|---|---|---|---|---|---|
| light - med | $1.0043 \times 10^{-9}$ | $1.0043 \times 10^{-9}$ | $1.0043 \times 10^{-9}$ | $1.0043 \times 10^{-9}$ | $1.0043 \times 10^{-9}$ | $1.0043 \times 10^{-9}$ | $1.0043 \times 10^{-9}$ |
| light - hard | $1.0043 \times 10^{-9}$ | $1.0043 \times 10^{-9}$ | $1.0043 \times 10^{-9}$ | 0.0401 | $1.0043 \times 10^{-9}$ | $1.0043 \times 10^{-9}$ | $1.0043 \times 10^{-9}$ |
| med - hard | $1.0043 \times 10^{-9}$ | $1.0043 \times 10^{-9}$ | $1.0043 \times 10^{-9}$ | $1.0043 \times 10^{-9}$ | $1.0043 \times 10^{-9}$ | $1.0043 \times 10^{-9}$ | $1.0043 \times 10^{-9}$ |

B

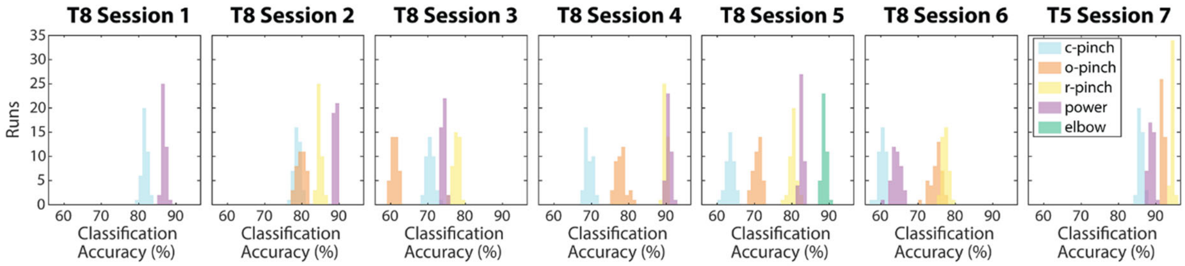


| | $p_1$ | $p_2$ | $p_3$ | $p_4$ | $p_5$ | $p_6$ | $p_7$ |
|---|---|---|---|---|---|---|---|
| cp - op | N/A | $5.0322 \times 10^{-7}$ | $9.0958 \times 10^{-9}$ | $9.0958 \times 10^{-9}$ | $1.7808 \times 10^{-8}$ | $9.0958 \times 10^{-9}$ | $9.0958 \times 10^{-9}$ |
| cp - rp | N/A | $9.0958 \times 10^{-9}$ | $9.0958 \times 10^{-9}$ | $9.0958 \times 10^{-9}$ | $1.7808 \times 10^{-8}$ | $9.0958 \times 10^{-9}$ | $9.0958 \times 10^{-9}$ |
| cp - pow | $7.4196 \times 10^{-9}$ | $9.0958 \times 10^{-9}$ | $9.0958 \times 10^{-9}$ | $9.0958 \times 10^{-9}$ | $1.7808 \times 10^{-8}$ | $9.0958 \times 10^{-9}$ | $9.0958 \times 10^{-9}$ |
| cp - elb | N/A | N/A | N/A | N/A | $1.7808 \times 10^{-8}$ | N/A | N/A |
| op - rp | N/A | $9.0958 \times 10^{-9}$ | $9.0958 \times 10^{-9}$ | $9.0958 \times 10^{-9}$ | $1.7808 \times 10^{-8}$ | $9.0958 \times 10^{-9}$ | $9.0958 \times 10^{-9}$ |
| op - pow | N/A | $9.0958 \times 10^{-9}$ | $9.0958 \times 10^{-9}$ | $9.0958 \times 10^{-9}$ | $1.7808 \times 10^{-8}$ | $9.0958 \times 10^{-9}$ | $9.0958 \times 10^{-9}$ |
| op - elb | N/A | N/A | N/A | N/A | $1.7808 \times 10^{-8}$ | N/A | N/A |
| rp - pow | N/A | $9.0958 \times 10^{-9}$ | $9.0958 \times 10^{-9}$ | $2.4722 \times 10^{-4}$ | $1.7808 \times 10^{-8}$ | $9.0958 \times 10^{-9}$ | $9.0958 \times 10^{-9}$ |
| rp - elb | N/A | N/A | N/A | N/A | $1.7808 \times 10^{-8}$ | N/A | N/A |
| pow - elb | N/A | N/A | N/A | N/A | $1.7808 \times 10^{-8}$ | N/A | N/A |

**Figure 7-1.** Statistics for go-phase force and grasp classifications accuracies. **A.** Force classification accuracy histograms (row 1) and corrected p values (row 2). Hard and light forces are classified significantly more accurately than medium forces across all sessions ($p < 0.05$). **B.** Grasp classification accuracy histograms (row 1) and corrected p values (row 2). Decoding performance differed significantly between grasps across all sessions.

A

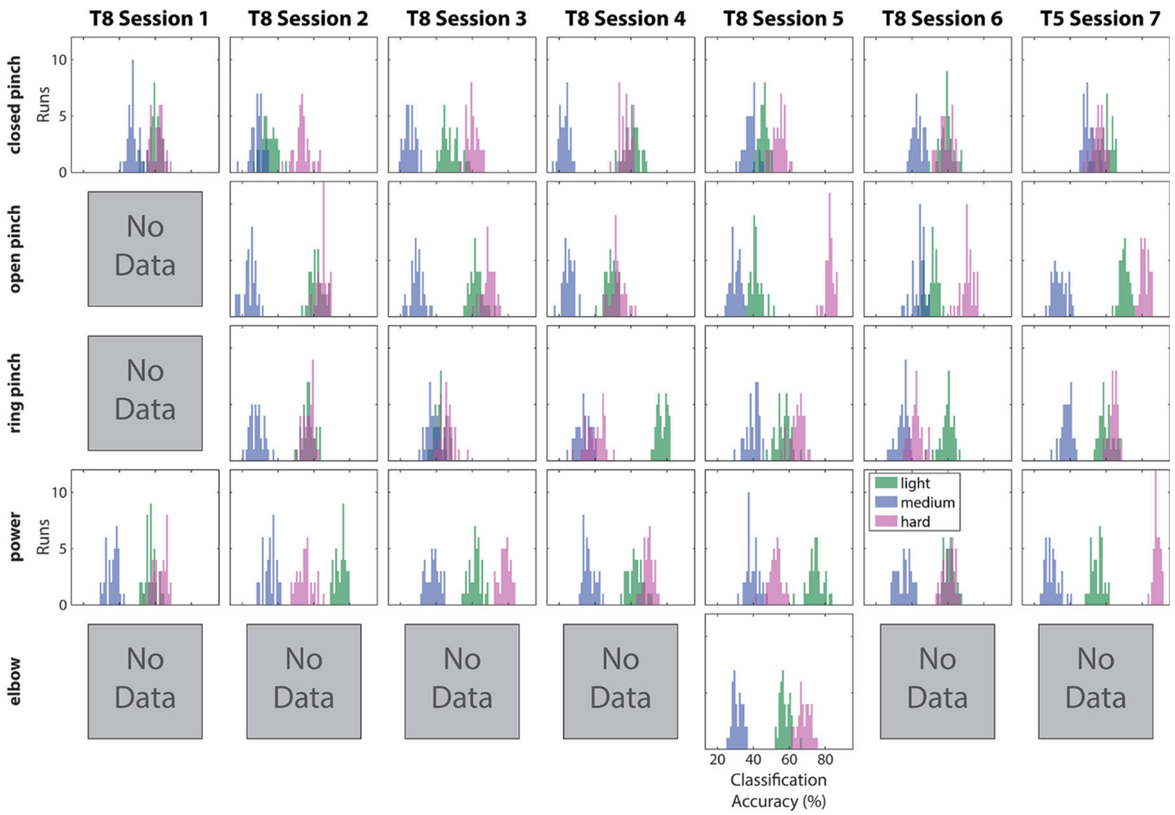


B

| | | $p_1$ | $p_2$ | $p_3$ | $p_4$ | $p_5$ | $p_6$ | $p_7$ |
|---|---|---|---|---|---|---|---|---|
| closed pinch | light - med | $1.5212 \times 10^{-9}$ | $3.4549 \times 10^{-9}$ | $1.5212 \times 10^{-9}$ | $1.5212 \times 10^{-9}$ | $1.5212 \times 10^{-9}$ | $1.5212 \times 10^{-9}$ | $1.5212 \times 10^{-9}$ |
| | light - hard | 0.2136 | $1.5212 \times 10^{-9}$ | $1.5212 \times 10^{-9}$ | $2.8060 \times 10^{-5}$ | $1.5212 \times 10^{-9}$ | 1.1250 | $4.2231 \times 10^{-5}$ |
| | med - hard | $1.5212 \times 10^{-9}$ | $1.5212 \times 10^{-9}$ | $1.5212 \times 10^{-9}$ | $1.5212 \times 10^{-9}$ | $1.5212 \times 10^{-9}$ | $1.5212 \times 10^{-9}$ | $1.6763 \times 10^{-9}$ |
| open pinch | light - med | N/A | $1.5212 \times 10^{-6}$ | $1.5212 \times 10^{-6}$ | $1.5212 \times 10^{-6}$ | $1.5212 \times 10^{-6}$ | $1.5212 \times 10^{-6}$ | $1.5212 \times 10^{-6}$ |
| | light - hard | | 0.1165 | $1.5212 \times 10^{-6}$ | 0.0078 | $1.5212 \times 10^{-6}$ | $1.5212 \times 10^{-6}$ | $1.5212 \times 10^{-6}$ |
| | med - hard | | $1.5212 \times 10^{-6}$ | $1.5212 \times 10^{-6}$ | $1.5212 \times 10^{-6}$ | $1.5212 \times 10^{-6}$ | $1.5212 \times 10^{-6}$ | $1.5212 \times 10^{-6}$ |
| ring pinch | light - med | N/A | $1.5212 \times 10^{-9}$ | $6.6714 \times 10^{-7}$ | $1.5212 \times 10^{-9}$ | $1.5212 \times 10^{-9}$ | $1.5212 \times 10^{-9}$ | $1.5212 \times 10^{-9}$ |
| | light - hard | | 1.2912 | $2.4214 \times 10^{-4}$ | $1.5212 \times 10^{-9}$ | $1.5212 \times 10^{-9}$ | $1.5212 \times 10^{-9}$ | $2.1445 \times 10^{-7}$ |
| | med - hard | | $1.5212 \times 10^{-9}$ | $1.5212 \times 10^{-9}$ | $1.5726 \times 10^{-9}$ | $1.5212 \times 10^{-9}$ | $1.5212 \times 10^{-9}$ | $1.5212 \times 10^{-9}$ |
| power | light - med | $1.5212 \times 10^{-9}$ | $1.5212 \times 10^{-9}$ | $1.5212 \times 10^{-9}$ | $1.5212 \times 10^{-9}$ | $1.5212 \times 10^{-9}$ | $1.5212 \times 10^{-9}$ | $1.5212 \times 10^{-9}$ |
| | light - hard | $4.3855 \times 10^{-9}$ | $1.5212 \times 10^{-9}$ | $1.5212 \times 10^{-9}$ | $1.5212 \times 10^{-9}$ | $1.5212 \times 10^{-9}$ | 1.2218 | $1.5212 \times 10^{-9}$ |
| | med - hard | $1.5212 \times 10^{-9}$ | $1.5726 \times 10^{-9}$ | $1.5726 \times 10^{-9}$ | $1.5726 \times 10^{-9}$ | $1.5726 \times 10^{-9}$ | $1.5726 \times 10^{-9}$ | $1.5726 \times 10^{-9}$ |
| elbow | light - med | N/A | N/A | N/A | N/A | $1.5212 \times 10^{-9}$ | N/A | N/A |
| | light - hard | | | | | $1.5212 \times 10^{-9}$ | | |
| | med - hard | | | | | $1.5726 \times 10^{-9}$ | | |

**Figure 7-2.** Statistics for go-phase force classification accuracies within individual grasp types. A one-way ANOVA was implemented on force classification accuracies achieved during different grasp types. **A.** Force classification accuracy histograms. **B.** P values between force pairs, corrected for multiple comparisons across grasps and sessions using the Benjamini-Hochberg procedure. Within each grasp, hard and light forces were classified more accurately than medium forces across all sessions ($p < 0.05$).

A

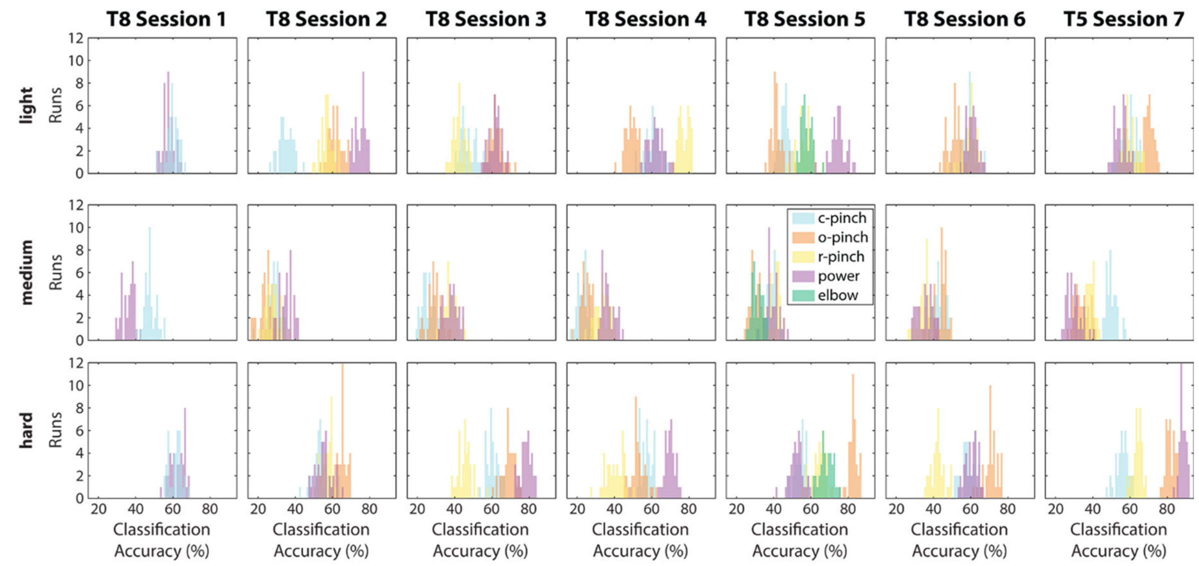


B

| | | $p_1$ | $p_2$ | $p_3$ | $p_4$ | $p_5$ | $p_6$ | $p_7$ |
|---|---|---|---|---|---|---|---|---|
| light | cp - op | N/A | $1.2174 \times 10^{-8}$ | $1.2174 \times 10^{-8}$ | $1.2174 \times 10^{-8}$ | $6.7371 \times 10^{-7}$ | $1.2174 \times 10^{-8}$ | $1.2174 \times 10^{-8}$ |
| | cp - rp | N/A | $1.2174 \times 10^{-8}$ | $1.3349 \times 10^{-8}$ | $1.2174 \times 10^{-8}$ | $2.3411 \times 10^{-8}$ | 1.6906 | 1.6099 |
| | cp - pow | $3.5015 \times 10^{-3}$ | $1.2174 \times 10^{-8}$ | $1.2174 \times 10^{-8}$ | 0.2546 | $2.3411 \times 10^{-8}$ | 0.5217 | $2.3930 \times 10^{-7}$ |
| | cp - elb | N/A | N/A | N/A | N/A | $2.3411 \times 10^{-8}$ | N/A | N/A |
| | op - rp | N/A | $8.0871 \times 10^{-8}$ | $1.2174 \times 10^{-8}$ | $1.2174 \times 10^{-8}$ | $2.3411 \times 10^{-8}$ | $1.2174 \times 10^{-8}$ | $1.2174 \times 10^{-8}$ |
| | op - pow | N/A | $1.2174 \times 10^{-8}$ | 1.5857 | $1.2174 \times 10^{-8}$ | $2.3411 \times 10^{-8}$ | $1.2174 \times 10^{-8}$ | $1.2174 \times 10^{-8}$ |
| | op - elb | N/A | N/A | N/A | N/A | $2.3411 \times 10^{-8}$ | N/A | N/A |
| | rp - pow | N/A | $1.2174 \times 10^{-8}$ | $1.2174 \times 10^{-8}$ | $1.2174 \times 10^{-8}$ | $2.3411 \times 10^{-8}$ | 0.8263 | $1.8119 \times 10^{-8}$ |
| | rp - elb | N/A | N/A | N/A | N/A | 1.5338 | N/A | N/A |
| | pow - elb | N/A | N/A | N/A | N/A | $2.3411 \times 10^{-8}$ | N/A | N/A |
| medium | cp - op | N/A | $6.5531 \times 10^{-8}$ | $4.1678 \times 10^{-6}$ | 0.0162 | $2.3411 \times 10^{-8}$ | 1.5773 | $1.2174 \times 10^{-8}$ |
| | cp - rp | N/A | 1.3853 | $1.2174 \times 10^{-8}$ | $1.2174 \times 10^{-8}$ | 0.4125 | $1.2174 \times 10^{-8}$ | $1.2174 \times 10^{-8}$ |
| | cp - pow | $1.2174 \times 10^{-8}$ | $1.2174 \times 10^{-8}$ | $1.2174 \times 10^{-8}$ | $1.2174 \times 10^{-8}$ | 0.8696 | $1.2174 \times 10^{-8}$ | $1.2174 \times 10^{-8}$ |
| | cp - elb | N/A | N/A | N/A | N/A | $2.3411 \times 10^{-8}$ | N/A | N/A |
| | op - rp | N/A | $1.1626 \times 10^{-5}$ | $1.2174 \times 10^{-8}$ | $1.2174 \times 10^{-8}$ | $2.3411 \times 10^{-8}$ | $1.2174 \times 10^{-8}$ | $4.4490 \times 10^{-6}$ |
| | op - pow | N/A | $1.2174 \times 10^{-8}$ | $1.2174 \times 10^{-8}$ | $1.2174 \times 10^{-8}$ | $2.3411 \times 10^{-8}$ | $1.2174 \times 10^{-8}$ | $4.2882 \times 10^{-7}$ |
| | op - elb | N/A | N/A | N/A | N/A | 1.0065 | N/A | N/A |
| | rp - pow | N/A | $1.2174 \times 10^{-8}$ | 1.6906 | $2.2072 \times 10^{-4}$ | 1.6916 | 1.3048 | $1.2174 \times 10^{-8}$ |
| | rp - elb | N/A | N/A | N/A | N/A | $2.3411 \times 10^{-8}$ | N/A | N/A |
| | pow - elb | N/A | N/A | N/A | N/A | $2.3411 \times 10^{-8}$ | N/A | N/A |
| hard | cp - op | N/A | $1.2174 \times 10^{-8}$ | $1.2174 \times 10^{-8}$ | $2.5703 \times 10^{-6}$ | $2.3411 \times 10^{-8}$ | $1.2174 \times 10^{-8}$ | $1.2174 \times 10^{-8}$ |
| | cp - rp | N/A | 0.0010 | $1.2174 \times 10^{-8}$ | $1.2174 \times 10^{-8}$ | $2.3411 \times 10^{-8}$ | $1.2174 \times 10^{-8}$ | $1.2174 \times 10^{-8}$ |
| | cp - pow | 0.1580 | 0.7988 | $1.2174 \times 10^{-8}$ | $1.2174 \times 10^{-8}$ | 0.1062 | 0.4024 | $1.2174 \times 10^{-8}$ |
| | cp - elb | N/A | N/A | N/A | N/A | $2.3411 \times 10^{-8}$ | N/A | N/A |
| | op - rp | N/A | $1.2174 \times 10^{-8}$ | $1.2174 \times 10^{-8}$ | $1.2174 \times 10^{-8}$ | $2.3411 \times 10^{-8}$ | $1.2174 \times 10^{-8}$ | $1.2174 \times 10^{-8}$ |
| | op - pow | N/A | $1.2174 \times 10^{-8}$ | $1.2174 \times 10^{-8}$ | $1.2174 \times 10^{-8}$ | $2.3411 \times 10^{-8}$ | $1.2174 \times 10^{-8}$ | $1.2174 \times 10^{-8}$ |
| | op - elb | N/A | N/A | N/A | N/A | $2.3411 \times 10^{-8}$ | N/A | N/A |
| | rp - pow | N/A | 0.1537 | $9.0958 \times 10^{-9}$ | $1.2174 \times 10^{-8}$ | $2.3411 \times 10^{-8}$ | $1.2174 \times 10^{-8}$ | $1.2174 \times 10^{-8}$ |
| | rp - elb | N/A | N/A | N/A | N/A | $1.0113 \times 10^{-5}$ | N/A | N/A |
| | pow - elb | N/A | N/A | N/A | N/A | $2.3411 \times 10^{-8}$ | N/A | N/A |

**Figure 7-3.** Statistics for go-phase force classification accuracies within individual force levels. A one-way ANOVA was implemented on the force classification accuracies achieved during different grasp types. **A.** Force classification accuracy histograms, color-coded by the grasp type used to produce each force level. **B.** P values between pairs of grasps used to produce each individual force level, corrected for multiple comparisons across forces and sessions using the Benjamini-Hochberg procedure. The decoding performance for each discrete force level

was significantly different across grasps ($p < 0.05$), indicating that grasp type affected force decoding performance.
